# Epigenetic–splicing regulation of *hTERT* mediated by *hTAPAS*

**DOI:** 10.64898/2026.05.08.723733

**Authors:** Valentine Sarah Jane Patterson, Alicia Gina Limbach, Julia Aylin Dogruyol, Christiane Krämer, Lars Erichsen

## Abstract

Telomerase activity is primarily controlled by expression of the catalytic subunit *hTERT*, yet the mechanisms coordinating its transcriptional and post-transcriptional regulation remain incompletely understood. Paradoxically, hypermethylation of a CpG-rich region upstream of the *hTERT* transcription start site, termed the *TERT* Hypermethylated Oncological Region (THOR), is strongly associated with *hTERT* activation. Recent studies suggest that THOR methylation influences the expression of *hTAPAS*, a long non-coding RNA transcribed antisense to the *hTERT* promoter that represses *hTERT* expression.

Here, we investigated the relationship between DNA methylation and alternative splicing of *hTERT*. Using CRISPR–Cas9–mediated targeted enrichment combined with Nanopore sequencing, we generated high-resolution DNA methylation maps across several kilobases surrounding the *hTERT* promoter and intronic regions encompassing exons 6–8 in multiple human cell lines. These analyses revealed distinct methylation signatures that correlate with specific *hTERT* splice isoform profiles.

Functional perturbation experiments further demonstrated that altering the epigenetic state of the locus influences *hTERT* splicing. Overexpression of the antisense lncRNA *hTAPAS*, as well as treatment with the DNA demethylating agent 5′-Azacytidine, reduced CpG methylation at the *hTERT* promoter and induced a shift in *hTERT* splice isoform distribution.

Together, our findings identify DNA methylation as an upstream regulator of *hTERT* alternative splicing and implicate the antisense lncRNA *hTAPAS* in shaping this regulatory landscape. The results point to an epigenetic mechanism linking *hTAPAS* to *hTERT* promoter methylation and splice isoform selection at the *hTERT* locus and provide new insight into how telomerase regulation may be remodeled during development and in cancer.

## Introduction

Telomerase maintains chromosome ends and controls cellular proliferative capacity through the catalytic subunit *hTERT*[1]. In humans and other vertebrates, telomeres consist of GT-rich (TTAGGG)_n_ repeats that cap chromosomal ends[2, 3]. *hTERT* expression is typically low in somatic cells with estimated fewer than five copies per cell and correlates closely with telomerase enzymatic activity[4]. During embryogenesis, telomerase activity is tightly regulated: embryonic stem cells exhibit high activity, which is largely extinguished during differentiation in most somatic lineages[5]. Conversely, telomerase reactivation is observed in approximately 90% of human cancers, regardless of tissue origin[6], suggesting a critical and temporally precise role for *hTERT* reexpression in carcinogenesis[7, 8].

Paradoxically, dense methylation of a CpG-rich region upstream of the *hTERT* transcription start site, termed the *hTERT* Hypermethylated Oncological Region (THOR), is strongly associated with *hTERT* activation[9]. This contradicts the canonical view that promoter methylation suppresses transcription, suggesting the existence of noncanonical epigenetic mechanisms regulating *hTERT.* The THOR region is located upstream of the core promoter and encompasses 433 bp with 52 CpG sites (chr5:1,295,321–1,295,753, GRCh37/hg19). In addition to THOR hypermethylation, recurrent *hTERT* promoter mutations create de novo ETS transcription factor binding sites that drive *hTERT* transcription in many cancers, highlighting multiple mechanisms by which telomerase can be reactivated during tumorigenesis[10].

Emerging evidence implicates a 1.6-kb long non-coding RNA transcribed antisense to the *hTERT* promoter (*hTAPAS*) in this regulatory axis[11, 12]. *hTAPAS* expression is inversely correlated with *hTERT* levels, and overexpression of *hTAPAS* suppresses *hTERT*, indicating that antisense transcription can modulate telomerase expression[11,12]. Furthermore, chromatin accessibility and histone modifications at the *hTERT* promoter contribute to transcriptional regulation, with permissive marks facilitating transcription factor binding and reactivation in cancer cells[14]. Alternative splicing provides an additional layer of *hTERT* regulation. The *hTERT* gene, spanning 42 kb and comprising 16 exons, has been reported to yield at least 21 splice variants[15, 16]. Apart from the full-length (FL) transcript incorporating all 16 exons, none of these alternatively spliced forms possesses reverse transcriptase activity, nor are they capable of telomere elongation[17, 18]. The most prevalent *hTERT* splice isoforms are the minus-alpha (α⁻), minus-beta (β⁻), and combined minus-alpha-beta (α⁻β⁻) variants all disrupting the reverse transcriptase domain and yield inactive proteins[19]. The balance between catalytically active full-length *hTERT* and inactive splice isoforms therefore represents a critical regulatory checkpoint for telomerase activity in both development and cancer. Splicing decisions are influenced by cell type and epigenetic context, yet the upstream determinants linking DNA methylation, lncRNA activity, and isoform selection remain poorly defined.

A growing body of evidence links DNA methylation to splicing regulation. In 2005, Young et al. demonstrated that methylated exonic DNA recruits MeCP2, which interacts with splicing factors such as YB-1[20]. Later, Shukla et al. showed that methylation-sensitive binding of CTCF at exon 5 of the human *CD45* gene regulates RNA polymerase II pausing and exon inclusion[21]. Together, these findings suggest that DNA methylation not only regulates transcription initiation but also influences co-transcriptional splicing decisions, a mechanism increasingly recognized as important for telomerase activity.

Here, we tested whether *hTAPAS* mediates the link between THOR methylation and *hTERT* splicing by using CRISPR–Cas9–based targeted enrichment combined with Nanopore sequencing to map DNA methylation across the *hTERT* promoter and introns 6–8. We identified methylation signatures that correlate with splice isoform usage. Furthermore, we demonstrate that perturbing the locus through *hTAPAS* overexpression or 5′-Azacytidine treatment shifts *hTERT* splicing patterns. These findings point to an epigenetic–splicing regulatory axis at *hTERT*, linking promoter methylation and *hTAPAS* antisense transcription to isoform-specific telomerase control.

## Results

### Inverse expression relationship between *hTAPAS* and *hTERT*

Previous reports suggest that the antisense lncRNA *hTAPAS* negatively regulates *hTERT* expression[11–12]. To characterize the relationship between *hTERT* expression, *hTAPAS* lncRNA levels and *hTERT* splice isoform usage, we analyzed a panel of human cell types compromising pluripotent (iPSC), immortalized (HEK293T), telomerase negative (VA13), primary fibroblasts (HFF-1, IMR90, BJ) and hematopoietic cells (NALM6) (Fig. 1A–B). Quantitative qRT-PCR analysis revealed pronounced cell type-specific differences in *hTERT* expression. Consistent with prior observations, an inverse correlation between *hTAPAS* and *hTERT* mRNA levels was evident in HFF-1 and BJ fibroblasts (*hTERT* absent, *hTAPAS* expressed) and in HEK293T and NALM6 cells (*hTERT* expressed, *hTAPAS* absent) (Fig. 1B). Intermediate patterns were observed in IMR90, VA13, and iPSCs: VA13 lacked *hTAPAS* with moderate *hTERT* expression, whereas IMR90 showed partial downregulation of *hTAPAS* with undetectable *hTERT* expression.

**Figure 1:**
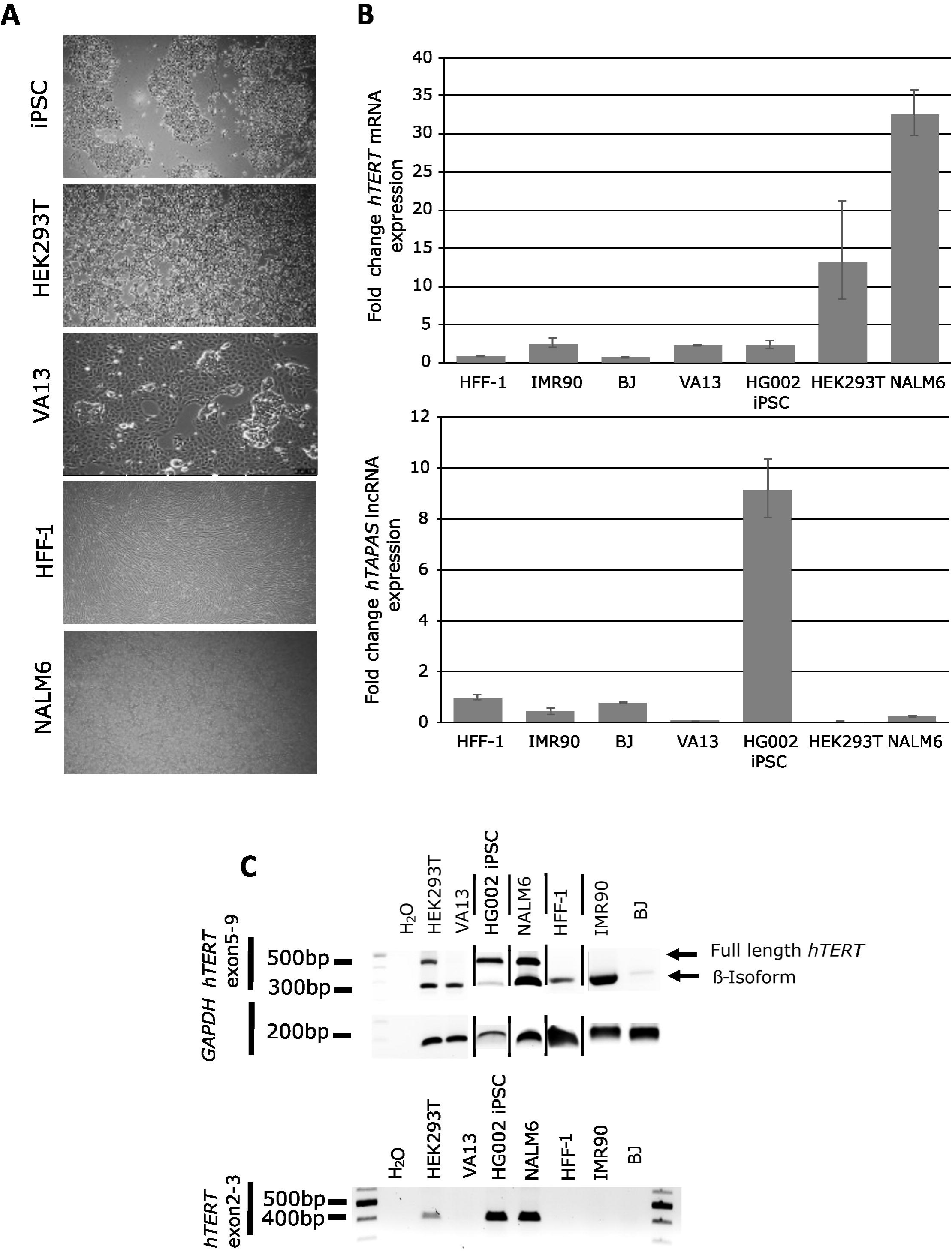
Relationship between *hTAPAS* and *hTERT* expression across human cell lines and associated splice isoforms. (A) BF images five out of the seven cell lines, including telomerase-negative fibroblasts (HFF-1, IMR90, BJ), telomerase-positive lines (HEK293T, NALM6, HG002 iPSCs), and the ALT-positive line VA13. (B) Quantification of *hTAPAS* and *hTERT* mRNA levels by qRT-PCR. (C) RT–PCR analysis of *hTERT* splice isoforms using primers spanning exons 2–3 and 5–9. An inverse relationship between *hTAPAS* and *hTERT* expression was observed in HFF-1 and BJ fibroblasts (low *hTERT*, detectable *hTAPAS*) and in HEK293T and NALM6 cells (high *hTERT*, absent *hTAPAS*), whereas intermediate patterns were detected in IMR90, VA13, and iPSCs, with iPSCs maintaining moderate *hTERT* expression despite high *hTAPAS* levels.

To further assess whether these differences are associated with changes in *hTERT* transcript composition, we examined alternative splicing by RT-PCR using primers spanning exon junctions 2–3 and 5–9. This analysis revealed that fibroblasts lack detectable full length *hTERT* mRNA expression. However, in HFF-1 and IMR90 cells, a band corresponding to the β isoform was observed (Fig. 1C). In contrast, HEK293T and NALM6 cells expressed both full-length and β isoforms. Induced pluripotent stem cells (iPSCs) predominantly expressed the full-length transcript, whereas VA13 cells exclusively expressed the β isoform. Exon 2–containing isoforms were restricted to cell lines dependent on *hTERT* activity for immortalization (HEK293T, NALM6 and HG002 iPSCs). The identity of amplification products was confirmed by Sanger sequencing (Supplementary Fig. 1). Collectively, these data demonstrate that *hTERT* expression, *hTAPAS* abundance, and *hTERT* splice isoform usage are tightly coordinated in a cell type-dependent manner. The observed inverse relationship between *hTAPAS* and *hTERT* expression, together with distinct splicing patterns, suggests a potential regulatory interplay between these factors.

### DNA methylation landscapes across the *hTERT* locus reveal cell type-specific heterogeneity

To investigate DNA methylation patterns across the *hTERT* locus, we performed CRISPR–Cas9-based targeted enrichment followed by Nanopore sequencing of a ∼9 kb region spanning *hTAPAS* through *hTERT* intron 2 (Chr. 5: 1,196,006–1,205,206) (Fig. 2) and a ∼6.5kb region within introns 6–8 (Chr. 5: 1,174,035-1,180,535) (Supplementary Fig. 2). Across all analyzed cell lines, DNA methylation patterns exhibited pronounced regional variability. While CpG methylation within intron 2 was consistently high (60–100%) local variability was observed at individual CpG sites across all samples. More heterogeneous methylation profiles among the studied cell lines were observed across *hTAPAS*, the THOR region, exon 2, and introns 6–8. CpGs within the *hTAPAS* region were nearly fully methylated in telomerase-positive lines (HEK293T, NALM6 and iPSC) and in the ALT-positive VA13, while largely unmethylated in fibroblasts except for partial methylation in IMR90 (∼50%). The core hTERT promoter and the region extending to exon 2 remained largely unmethylated in fibroblasts and HG002 iPSCs. In contrast, VA13 exhibited pronounced hypermethylation (60–100%), whereas NALM6 and HEK293T cells displayed intermediate methylation levels, with approximately 40% of CpG sites methylated. Methylation differences were most pronounced within exon 2. In all fibroblast lines, CpGs were completely unmethylated in the beginning of exon 2, beyond which methylation levels sharply increased to nearly 100%. In VA13, a gradual decline across exon 2 methylation was observed, from ∼60% to ∼20% across Exon. Conversely, exon 2 CpGs showed only low levels of methylation in HG002 iPSCs (∼20%), while HEK293T and NALM6 exhibited complete methylation across the entire exon.

**Figure 2:**
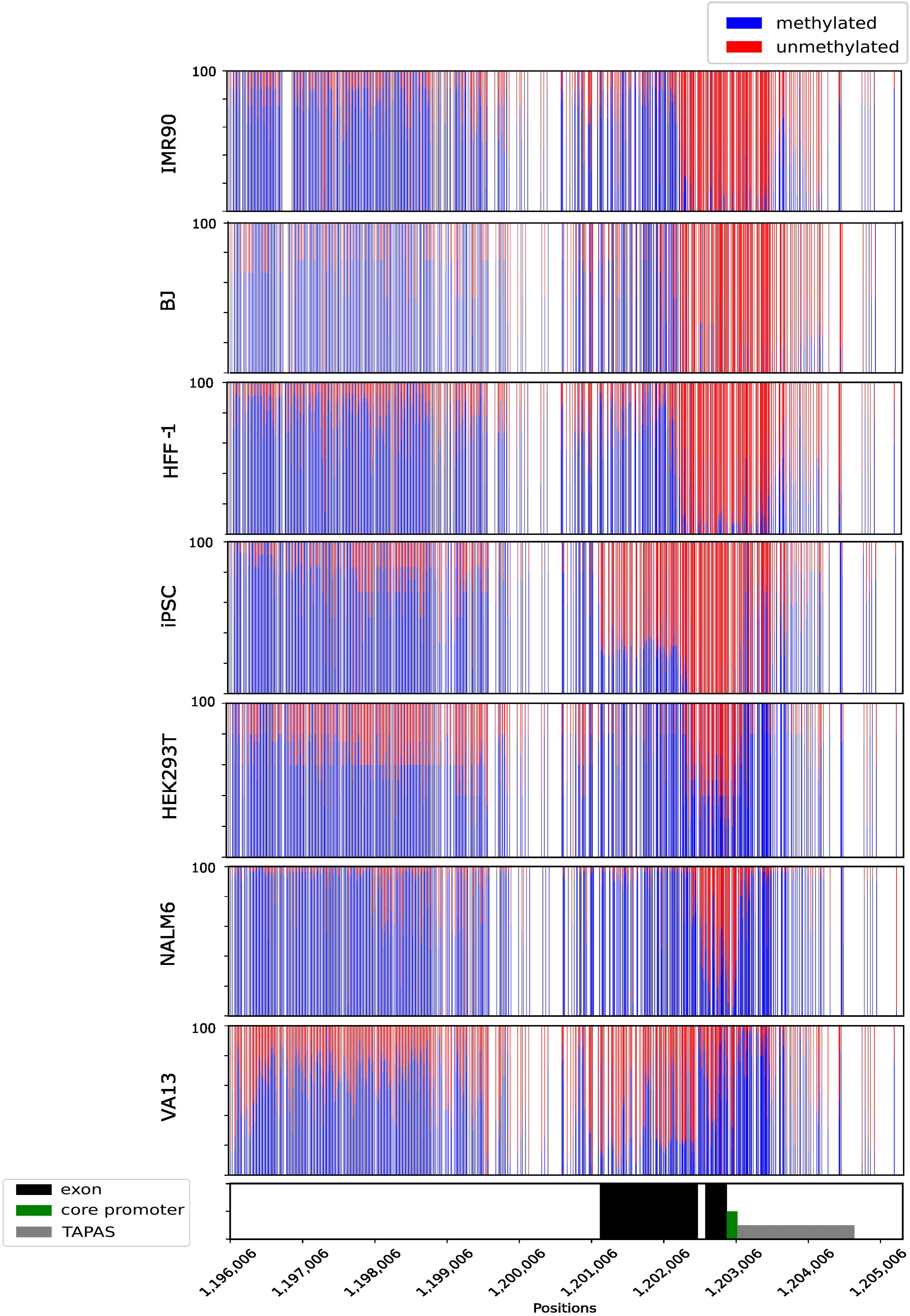
DNA methylation landscapes across the *hTERT* locus in distinct human cell lines. Targeted DNA methylation profiling was performed using CRISPR–Cas9 enrichment followed by Nanopore sequencing across a ∼9 kb region spanning *hTAPAS* through *hTERT* intron 2 (Chr. 5: 1,196,006–1,205,206) and a ∼6.5 kb region covering introns 6–8 (Chr. 5: 1,174,035–1,180,535). The analysis included telomerase-negative fibroblasts (HFF-1, IMR90, BJ), telomerase-positive cell lines (HEK293T, NALM6, HG002 iPSCs), and the ALT-positive cell line VA13. DNA methylation levels at individual CpG sites are depicted across the indicated genomic regions, including *hTAPAS*, the THOR region, the core promoter, exon1, intron1, exon 2 and intron 2. Methylation for each individual CpG is shown as a percentage, with unmethylated CpGs depicted in red and methylated CpGs in blue. Methylation across intron 2 was consistently high (80–100%) in all cell lines, whereas regions encompassing *hTAPAS*, the THOR region, exon 2, and introns 6–8 displayed marked variability between cell types. CpGs within the *hTAPAS* region were highly methylated in telomerase-positive cells and in the ALT-positive VA13 line, but largely unmethylated in fibroblasts, with partial methylation observed in IMR90. The core *hTERT* promoter and exon 2–proximal regions remained mostly unmethylated in all cell lines except VA13, which exhibited substantial hypermethylation

Within introns 6-8 (Supplementary Fig. 2), methylation patterns were similarly heterogeneous and cell type specific. In telomerase-positive HEK293T cells and iPSCs a near-complete CpG methylation was observed (∼90-100%). In contrast VA13 cells showed a highly heterogeneous methylation landscape. Methylation levels were 40–60% across the variable number of tandem repeats (VNTR) region upstream of exon 7, decreased sharply to ∼20% between exons 7 and 8, and then rose again to nearly 90% within a ∼500 bp region downstream of exon 8, before declining to ∼20% across the remainder of intron 8.

Together, these results demonstrate that DNA methylation patterns across the *hTERT* locus is highly region-specific and differs substantially between cell lines. However, the observed patterns do not follow a uniform or binary distribution but rather reflect a complex and heterogeneous epigenetic landscape.

### *hTAPAS* overexpression alters *hTERT* expression, splice isoform distribution, and DNA methylation patterns

The THOR region has recently been proposed to function as the primary regulatory unit of *hTAPAS*, rather than *hTERT*, as hypermethylation of this region has been associated with repression of *hTAPAS* and concomitant activation of *hTERT*[12]. To investigate whether *hTAPAS* influences DNA methylation dynamics in trans, we cloned the full-length *hTAPAS* transcript into a pcDNA-3xHA expression vector containing an *eGFP* reporter. To distinguish sequence-specific effects from general RNA-related or transcription-dependent effects, we additionally generated an antisense construct of *hTAPAS*, hereafter referred to as *hSAPAT*. Both constructs, together with the empty pcDNA-3xHA vector as a control, were transiently transfected into HEK293T and VA13 cells.

Transfection efficiency was confirmed by monitoring *eGFP* expression using fluorescence microscopy and fluorescence-activated cell sorting (FACS) (Supplementary Fig. 4+5). Proliferation assays were performed over a 144-hour course using a Nexcelom automated cell counter (Fig. 3A and Fig. 4A). In both cell lines, expression of *hSAPAT* resulted in only a minor, non-significant reduction in cell number compared with control conditions at 24, 72, and 144 hours post transfection. In contrast, expression of *hTAPAS* led to a pronounced and statistically significant decrease in cell number relative to both the empty vector control and the *hSAPAT* condition.

**Figure 3:**
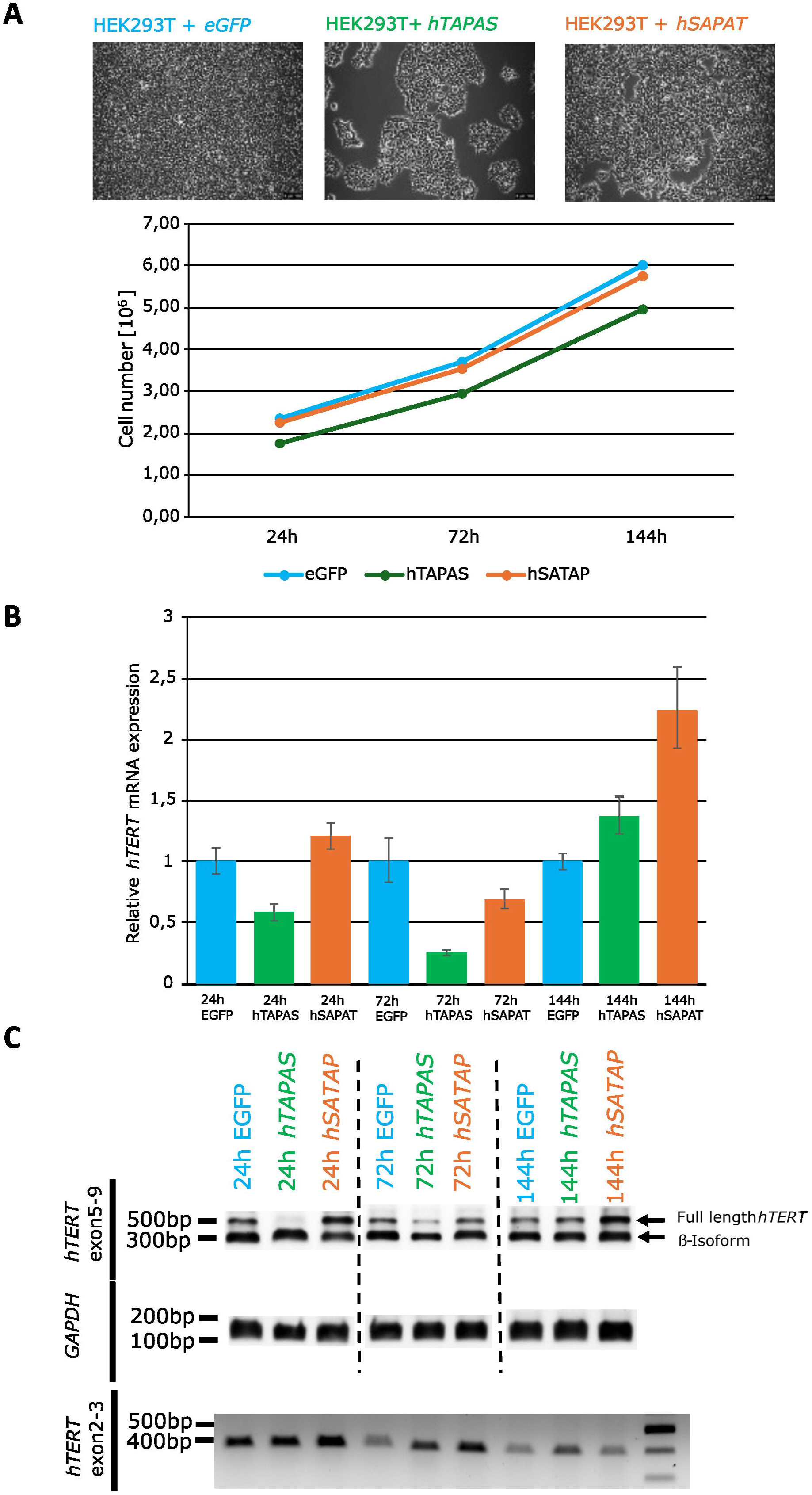
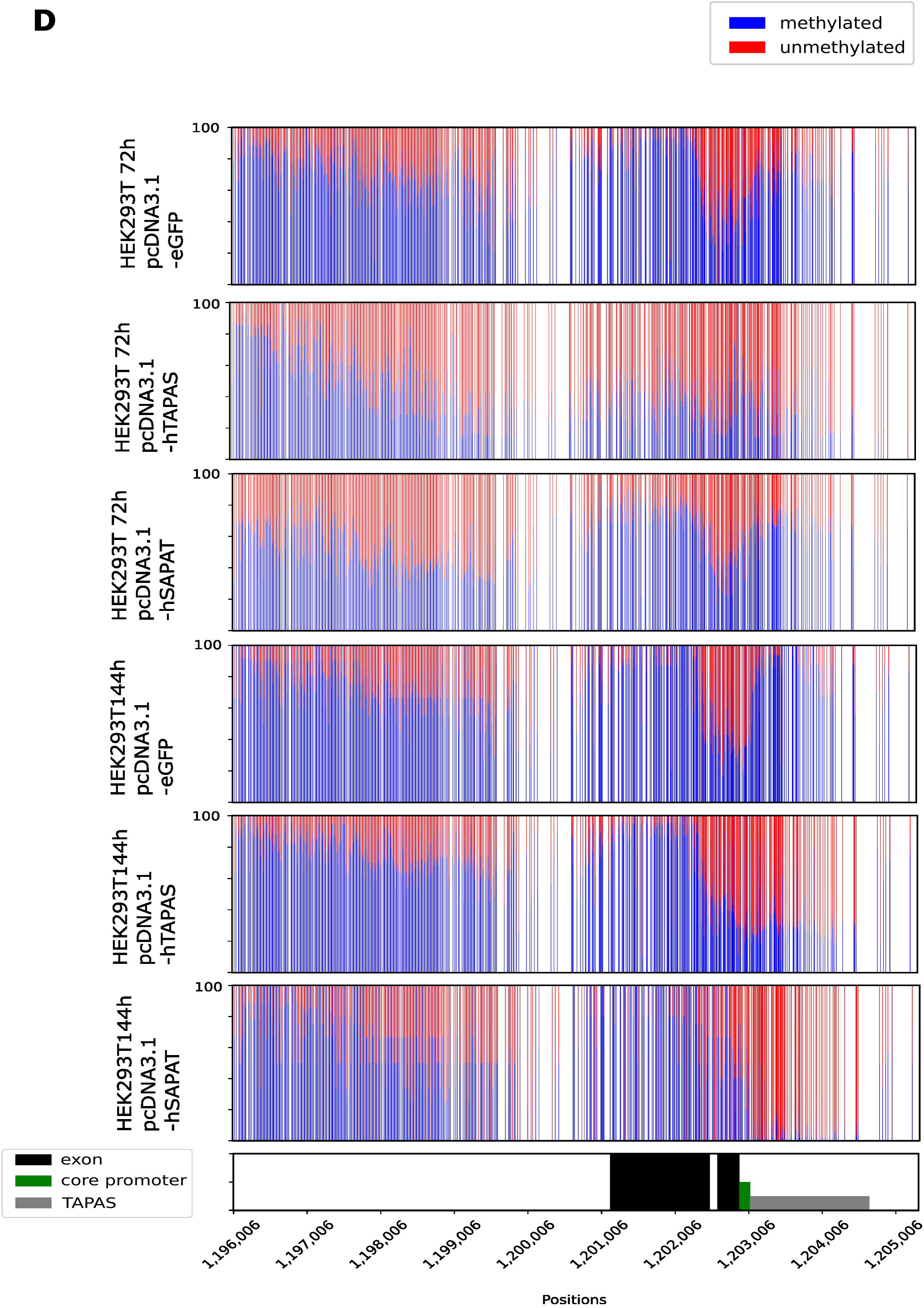
Effects of ectopic *hTAPAS* overexpression on *hTERT* expression, alternative splicing, and DNA methylation in HEK293T cells. HEK293T cells were transiently transfected with pcDNA3.1-eGFP (empty vector), full-length *hTAPAS*, or antisense *hTAPAS* (*hSAPAT*) to assess effects on cell growth, *hTERT* regulation, and local DNA methylation. (A) BF images and proliferation over 144 hours post-transfection was quantified using an automated cell counter. *hTAPAS* expression significantly reduced cell numbers compared with empty vector and *hSAPAT* controls, whereas *hSAPAT* had minor, non-significant effects. (B) Quantification of *hTAPAS* and *hTERT* mRNA levels by qRT-PCR. (C) RT–PCR analysis of *hTERT* splice isoforms using primers spanning exons 2–3 and 5–9. The full-length *hTERT* isoform was abolished and non-functional variants predominated at early time points. Splicing patterns largely returned to baseline by 144 hours. (D) DNA methylation dynamics across the ∼9 kb region spanning *hTAPAS* through *hTERT* intron 2 (Chr. 5: 1,196,006–1,205,206) are shown as percentages per CpG site, with unmethylated CpGs in red and methylated CpGs in blue. Hypomethylation extended from *hTAPAS* through exon 2. Differences between *hTAPAS* and *hSAPAT* were particularly across the *hTAPAS*–exon 2 segment, with *hTAPAS* inducing stronger hypomethylation (10–50% CpG methylation) compared with *hSAPAT* (70–80%). By 144 hours, methylation levels partially recovered, while the *hTAPAS* locus and THOR region remained strongly hypomethylated (2–10% CpG methylation).

**Figure 4:**
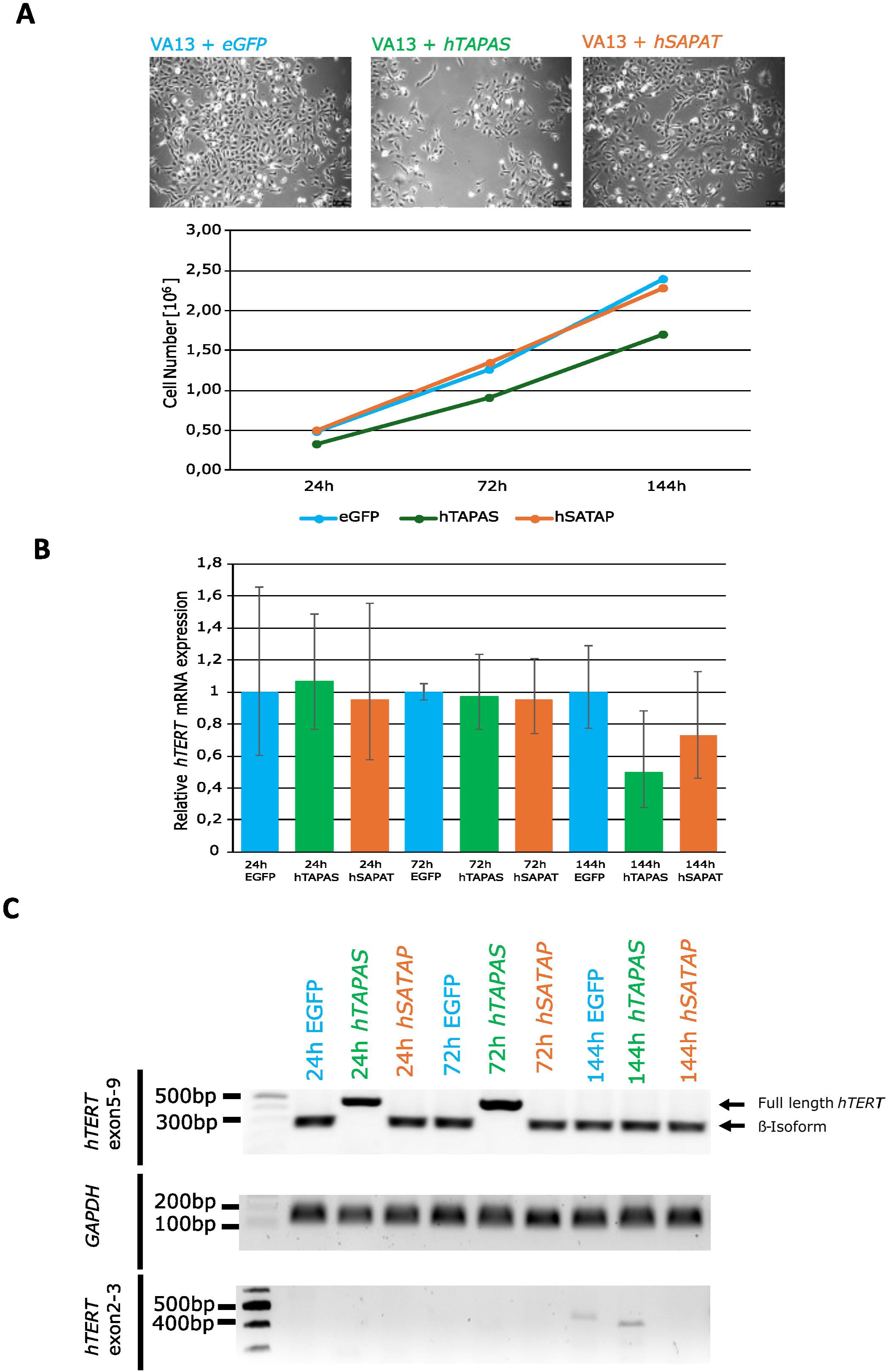
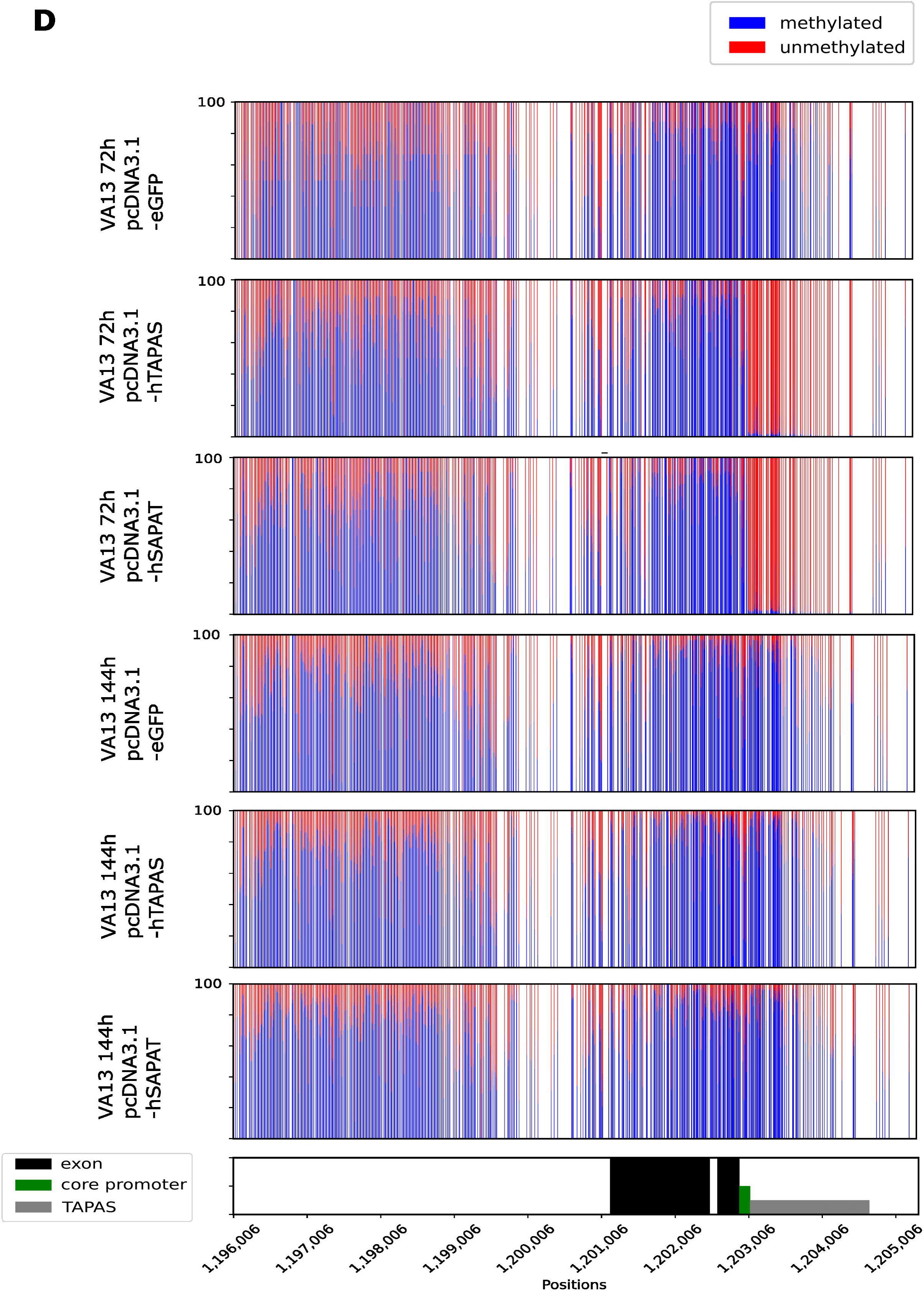
Effects of ectopic *hTAPAS* overexpression on *hTERT* expression, alternative splicing, and DNA methylation in VA13 cells. VA13 cells were transiently transfected with pcDNA3.1-eGFP (empty vector), full-length *hTAPAS*, or antisense *hTAPAS* (hSAPAT) to assess effects on cell growth, *hTERT* regulation, and local DNA methylation. (A) BF images and proliferation over 144 hours post-transfection was quantified using an automated cell counter. *hTAPAS* expression significantly reduced cell numbers compared with empty vector and *hSAPAT* controls, whereas *hSAPAT* had minor, non-significant effects. (B) Quantification of *hTAPAS* and *hTERT* mRNA levels by qRT-PCR. (C) RT–PCR analysis of *hTERT* splice isoforms using primers spanning exons 2–3 and 5–9. Full-length *hTERT* was preferentially expressed at early time points. Splicing patterns largely returned to baseline by 144 hours. (D) DNA methylation dynamics across the ∼9 kb region spanning *hTAPAS* through *hTERT* intron 2 (Chr. 5: 1,196,006–1,205,206) are shown as percentages per CpG site, with unmethylated CpGs in red and methylated CpGs in blue. Demethylation was largely restricted to *hTAPAS* through intron 1at 72h post transfection and recovered fully at 144h post transfection.

Quantitative real-time PCR confirmed robust expression of exogenous *hTAPAS* in transfected cells (Supplementary Fig. 6). Consistent with previous reports, *hTAPAS* overexpression resulted in downregulation of *hTERT* expression in both cell lines (Fig. 3B and Fig. 4B). In HEK293T cells, *hTERT* transcript levels were significantly reduced at 24 and 72 hours post transfection, followed by recovery to near-control levels at 144 hours. In VA13 cells, *hTERT* expression remained unchanged at early time points but was significantly reduced at 144 hours post transfection.

*hTAPAS* overexpression was associated with alterations in splice isoform usage in both cell lines (Figs. 3C and 4C). In HEK293T cells, *hTAPAS* expression abolished the full-length *hTERT* isoform and promoted the generation of non-functional splice variants at 24 and 72 hours post-transfection (Fig. 3C), whereas isoforms containing exon 2 appeared largely unchanged under these conditions. Conversely, in VA13 cells, *hTAPAS* overexpression induced a near-complete switch toward the full-length *hTERT* isoform at the same time points (Fig. 4C). Notably, in VA13 cells, a PCR product containing exon 2 was detectable only at 144 hours post-transfection with the *hTAPAS* construct. In both cell lines, canonical splicing patterns of exons 5-7 were restored 144 hours post-transfection.

Finally, we examined DNA methylation dynamics in HEK293T and VA13 cells 72 h and 144 h after transfection with pcDNA3.1-*eGFP*, *hTAPAS*, or *hSAPAT* expression constructs (Figs. 3D, 4D). At all-time points, the empty vector control exerted no detectable influence on the methylation landscape of the ∼9 kb interval encompassing *hTAPAS* through *hTERT* intron 2 (Chr. 5: 1,196,006–1,205,206). In marked contrast, overexpression of either *hTAPAS* or *hSAPAT* induced substantial DNA demethylation in both HEK293T and VA13 cells. In VA13 cells, this effect was confined to the *hTAPAS* region, which became almost completely unmethylated within 72 h of transfection. In HEK293T cells, by comparison, methylation levels were reduced across the entire examined interval at the same time point, declining to approximately 40–60% overall.

Whereas no appreciable differences between *hTAPAS* and *hSAPAT* were observed in VA13 cells, their effects diverged in HEK293T cells, most prominently within the segment spanning *hTAPAS* to exon 2. In *hSAPAT*-transfected cells, 70–80% of CpG sites within this region remained methylated. By contrast, *hTAPAS* overexpression resulted in markedly lower methylation levels (10–50%), with the highest residual methylation localized at the exon 1–exon 2 boundary. Notably, intron 2 exhibited a comparable reduction in methylation under both conditions, reaching ∼50% across the analysed region. By 144 h post-transfection, methylation levels had largely recovered, approaching near-complete CpG methylation in both cell lines and experimental conditions. In contrast, the *hTAPAS* locus, including the THOR region, remained strongly hypomethylated, with ∼10% CpG methylation following *hTAPAS* overexpression and only 2–5% after *hSAPAT* overexpression in HEK293T cells.

Collectively, these findings indicate that ectopic *hTAPAS* expression is associated with dynamic changes in *hTERT* expression, splice isoform distribution and local DNA methylation patterns. However, similar effects observed with the antisense construct suggest that at least part of the methylation changes may reflect RNA- or transcriptional associated effects rather than strictly sequence-specific mechanisms.

### DNA demethylation induces changes in *hTERT* expression, splicing and local methylation patterns

To determine the impact of DNA methylation on *hTERT* regulation, HEK293T and VA13 cells were treated with the DNA methyltransferase inhibitor 5’-Azacytidine. We exposed both cell lines to 1 µM 5’-Azacytidine for five days, observing pronounced cytotoxicity in HEK293T cells and moderate cell death in VA13 cells (Fig. 5A). This treatment reduced THOR methylation by 16–26% and induced locus-wide hypomethylation across *hTAPAS*, exon 1–2, and introns 6–8 (Supplementary Fig. 2+7, Fig. 5A+E). Residual methylation was retained at subsets of CpGs, indicating incomplete and heterogeneous demethylation.

**Figure 5:**
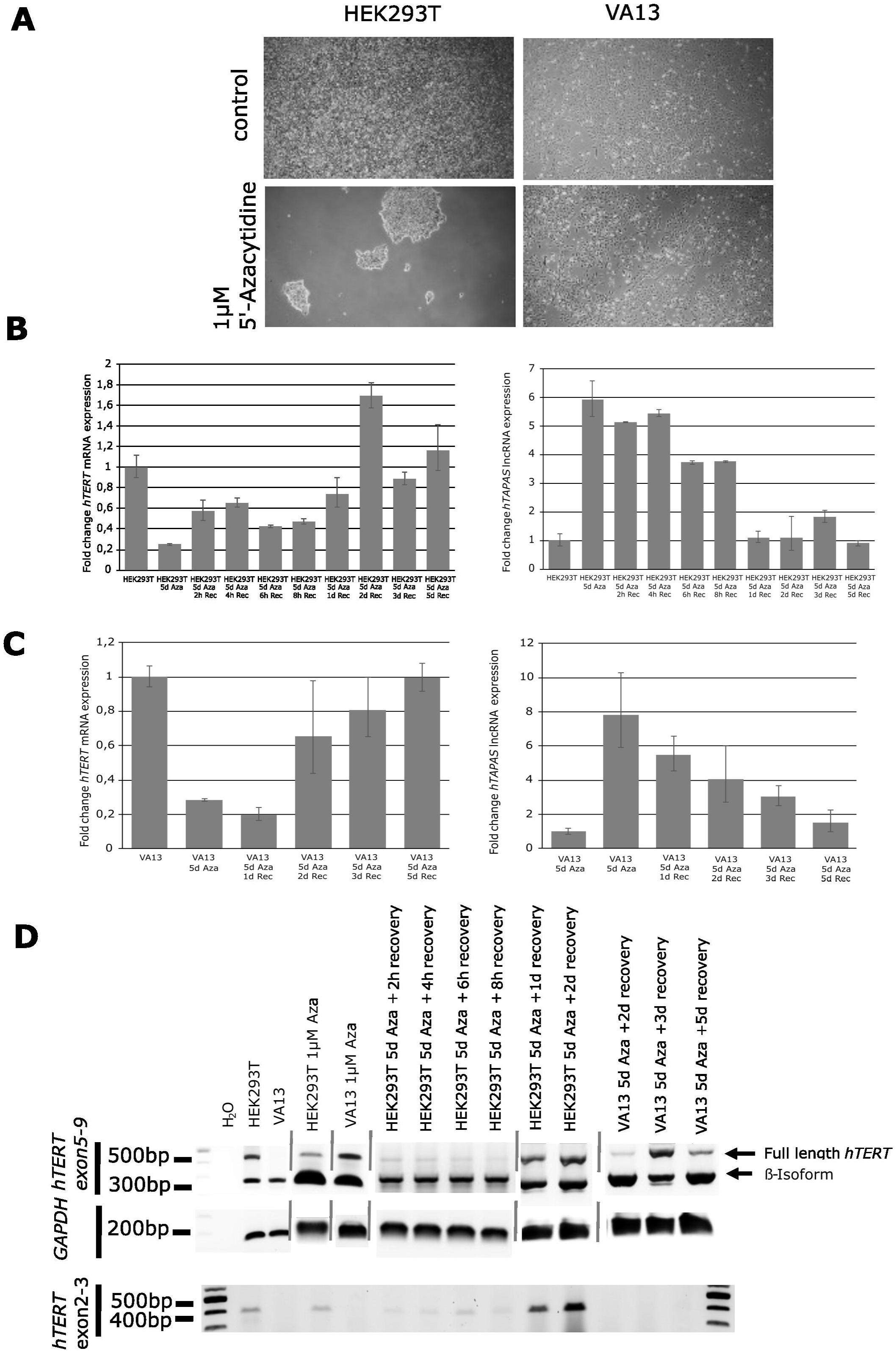
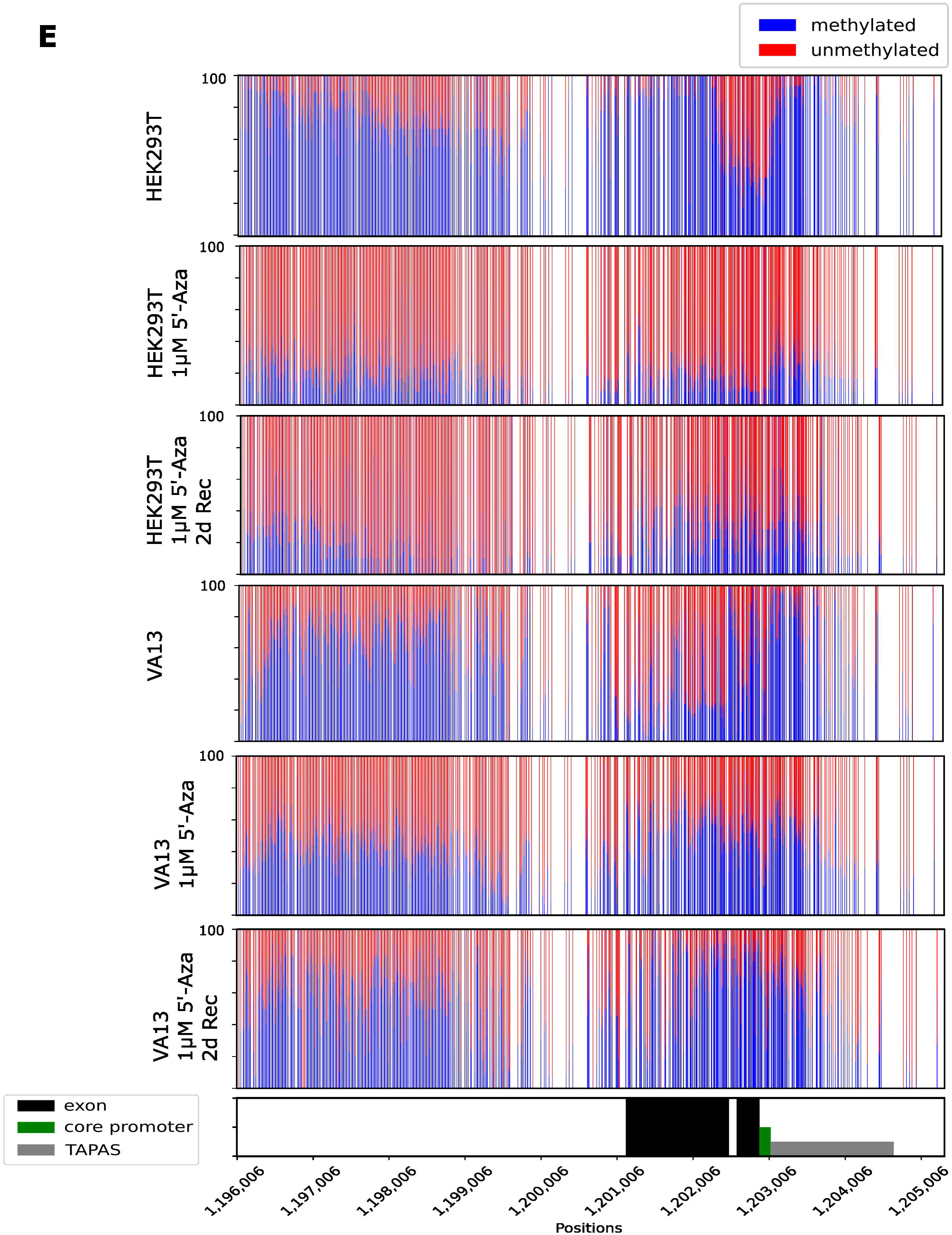
Demethylation-driven changes in *hTERT* expression and splicing. HEK293T and VA13 cells were treated with 1 µM 5′-Azacytidine for five days to inhibit DNA methyltransferase activity and induce locus-wide hypomethylation. (A) Cell viability over the treatment course showed pronounced cytotoxicity in HEK293T and moderate cell death in VA13 cells. (B–C) Quantification of *hTAPAS* and *hTERT* mRNA levels by qRT-PCR. Gene expression analysis demonstrated that demethylation decreased *hTERT* expression while reciprocally upregulating *hTAPAS*, consistent with their inverse relationship. (D) RT–PCR analysis of *hTERT* splice isoforms using primers spanning exons 2–3 and 5–9. RT–PCR analysis of *hTERT* splice isoforms showed a shift toward the β–isoform in HEK293T cells and induction of full-length *hTERT* in VA13 cells, mirroring the effects observed with *hTAPAS* overexpression. (E) DNA methylation dynamics across the ∼9 kb region spanning *hTAPAS* through *hTERT* intron 2 (Chr. 5: 1,196,006–1,205,206) are shown as percentages per CpG site, with unmethylated CpGs in red and methylated CpGs in blue. HEK293T cells rapidly remethylated the promoter–exon 2 region, while cells exon 2 CpGs got partially hypermethylated in VA13 cells upon 5’-Azacytidine treatment.

Demethylation resulted in decreased *hTERT* expression accompanied by reciprocal upregulation of *hTAPAS* in both cell lines (Figs. 5B, C). Concomitantly, splice isoform usage was found to be altered upon treatment. In HEK293T cells 5’-Azacytidine induced a shift toward the β-isoform, while in VA13 cells the induction of full-length *hTERT* isoforms was observed (Figs. 5D). However, in contrast to the *hTAPAS* overexpression experiments, 5′-Azacytidine treatment did not induce variants that lack exon 2 in HEK293T cells and failed to generate detectable exon 2–containing isoforms in VA13 cells.

Following withdrawal of 5′-Azacytidine, expression dynamics diverged between the two cell lines. In HEK293T cells, both *hTERT* and *hTAPAS* levels returned rapidly to baseline within 24 hours, whereas in VA13 cells, *hTERT* expression continued to decline transiently before gradually recovering over 120 hours, with *hTAPAS* displaying reciprocal kinetics. These transcriptional and splicing alterations closely paralleled the methylation dynamics of the promoter–exon 2 region in VA13 cells. In these cells, methylation at exon 2 CpG sites was maintained for several days, coinciding with sustained expression of full-length hTERT, even as distal intronic regions reverted to baseline methylation (Supplementary Figure 2). In contrast, HEK293T cells exhibited persistently low CpG methylation across the examined loci, although exon 2 showed a modest increase from 10% to 20% two days following 5′-Azacytidine withdrawal.

Together, these findings demonstrate that pharmacological DNA demethylation is associated with changes in *hTERT* expression and splice isoform usage, as well as dynamic and region-specific alterations in DNA methylation across the locus.

## Discussion

Our study reveals that DNA methylation, antisense transcription, and alternative splicing are integrated to orchestrate *hTERT* expression. Similar epigenetic–splicing coupling has been described at other loci. For example, DNA methylation–sensitive binding of CTCF regulates exon inclusion in the immune receptor gene PTPRC[21], while chromatin marks such as H3K36me3 control alternative exon usage in *FGFR2* through recruitment of splicing regulators[22]. Genome-wide analyses have further shown that intragenic DNA methylation can influence RNA polymerase II elongation through recruitment of methyl-binding proteins such as MeCP2, thereby shaping co-transcriptional splice-site recognition[20]. Our data indicate that the *hTAPAS*–exon 2 interval functions as a focal epigenetic hub in which methylation status shapes the local transcriptional environment and thereby biases splice-site selection, whereas methylation within introns 6–8 appears secondary to exon inclusion. This relationship was not clearly reflected in exon 2 inclusion itself, despite previous work[23] identifying exon 2 splicing as a key developmental switch governing *hTERT* expression. In that context, exon 2 inclusion has been linked to transcript stability and telomerase activity, whereas its exclusion promotes mRNA decay. Our findings suggest that, in the setting of *hTAPAS*-associated regulation, exon 2 usage may be modulated in a more context-dependent manner, rather than strictly following a binary switch-like behaviour. Our data point to three distinct, context-dependent epigenetic states:

1. Telomerase-negative, *hTAPAS*-positive fibroblasts displayed near-complete CpG methylation across the *hTERT* gene body, with the notable exception of the boundary between the *hTERT* core promoter and exon 1. In contrast, the *hTAPAS* locus remained largely unmethylated.
2. Telomerase-positive cancer cell lines exhibited a largely opposing profile, with pervasive methylation across both *hTAPAS* and *hTERT* while maintaining comparatively lower methylation at the promoter–exon 1 boundary.
3. Induced pluripotent stem cells (iPSCs) displayed a hybrid epigenetic configuration: extensive methylation across *hTAPAS* and intron 2 but pronounced hypomethylation from the core promoter through exon 2. This pattern, reminiscent of bivalent chromatin domains observed in pluripotent cells[24, 25], may permit concurrent expression of *hTAPAS* and *hTERT* while maintaining the capacity for rapid epigenetic remodelling during differentiation.

The divergent responses observed between HEK293T and VA13 cells further underscore the context dependence of this regulatory axis, as evidenced by differential DNA methylation patterns across exon 2 under varying cellular and experimental conditions. Moreover, these contrasting outcomes may likely reflect differences in chromatin architecture, transcriptional kinetics, and the availability or activity of splicing factors across distinct cellular environments. These factors warrant systematic investigation in future studies to delineate their precise contributions to the observed regulatory variability.

To establish these states, antisense transcription of *hTAPAS* appears not merely correlative but functionally determinant, placing it within a broader class of antisense transcripts that regulate gene expression through coordinated epigenetic and co-transcriptional mechanisms. *hTAPAS* expression is associated with the establishment and the maintenance of an unmethylated 5′-regulatory domain, thereby shaping local chromatin accessibility and coordinating splice-site selection. Hypermethylation of the TERT Hypermethylated Oncological Region (THOR) has been widely associated with activation of *hTERT* in human cancers, supporting the idea that epigenetic remodelling within the 5′-regulatory region of the locus plays a central role in telomerase reactivation[9]. Its overexpression recapitulates the effects of pharmacologic demethylation despite the well-known cell-cycle dependency and cytotoxicity of the nucleoside analogue 5′-Azacytidine[26, 27].

We propose that *hTAPAS* expression maintains an unmethylated 5′ regulatory domain extending through THOR and intron 2, potentially via recruitment of DNA demethylating machinery[28, 29]. In this feed-forward framework, loss of *hTAPAS* expression, driven by hypermethylation of its putative promoter during early carcinogenesis, would shift the locus toward a fully methylated state that favors the production of full-length, catalytically active *hTERT* isoforms.

Alternative splicing is tightly coupled with these epigenetic cues. The balance of full-length and inactive β⁻/α⁻ isoforms correlate with the methylation state of the *hTAPAS*–exon 2 interval, demonstrating that DNA methylation and antisense transcription jointly shape splice-site choice. Consistent with the importance of regulated splice-site selection in telomerase control, the splicing factor NOVA1 has been shown to promote inclusion of exons within the reverse transcriptase domain of *hTERT*, thereby favouring production of the catalytically active full-length transcript in cancer cells[30]. Recovery of DNA methylation and isoform distribution following withdrawal of 5′-Azacytidine or *hTAPAS* overexpression further underscores the locus’s epigenetic memory, likely maintained through replication-coupled DNMT1 activity. Together, these observations suggest that antisense transcription and DNA methylation cooperate to establish a chromatin environment that modulates RNA polymerase II elongation dynamics and thereby influences co-transcriptional splice-site recognition at the *hTERT* locus.

These findings also have broader implications for understanding how telomerase is reactivated during tumorigenesis. While promoter mutations[10, 13], chromatin remodelling[31, 32], and epigenetic activation[9] are known mechanisms that activate *hTERT* transcription, our results suggest that regulation of splice-site choice represents an additional layer controlling functional telomerase output. In this context, coordinated epigenetic regulation of antisense transcription and local DNA methylation may determine whether the locus produces catalytically active or inactive *hTERT* isoforms. Such a mechanism would allow cancer cells to tune telomerase activity without requiring large changes in overall transcriptional output. Targeting this regulatory axis, either through modulation of antisense transcription or locus-specific epigenetic editing, may provide a strategy to selectively shift *hTERT* splice isoform usage toward inactive variants.

Several limitations of the present study should be considered. Although our data demonstrate strong correlations between methylation patterns and *hTERT* splice isoform usage, the precise molecular intermediates linking DNA methylation to splice-site selection remain to be defined. Whether specific methyl-binding proteins or chromatin-associated splicing factors mediate this effect at the *hTERT* locus will require further investigation.

Collectively, these findings support a conceptual model (Fig. 6) in which coordinated methylation across *hTAPAS* and *hTERT* exon 2 functions as a central epigenetic switch controlling telomerase expression. In this framework, exon 2 emerges not only as a developmental checkpoint but also as an integration point for epigenetic and transcriptional inputs. This multilayered regulatory axis provides mechanistic insight into telomerase silencing during differentiation and its reactivation in tumorigenesis.

**Figure 6.**
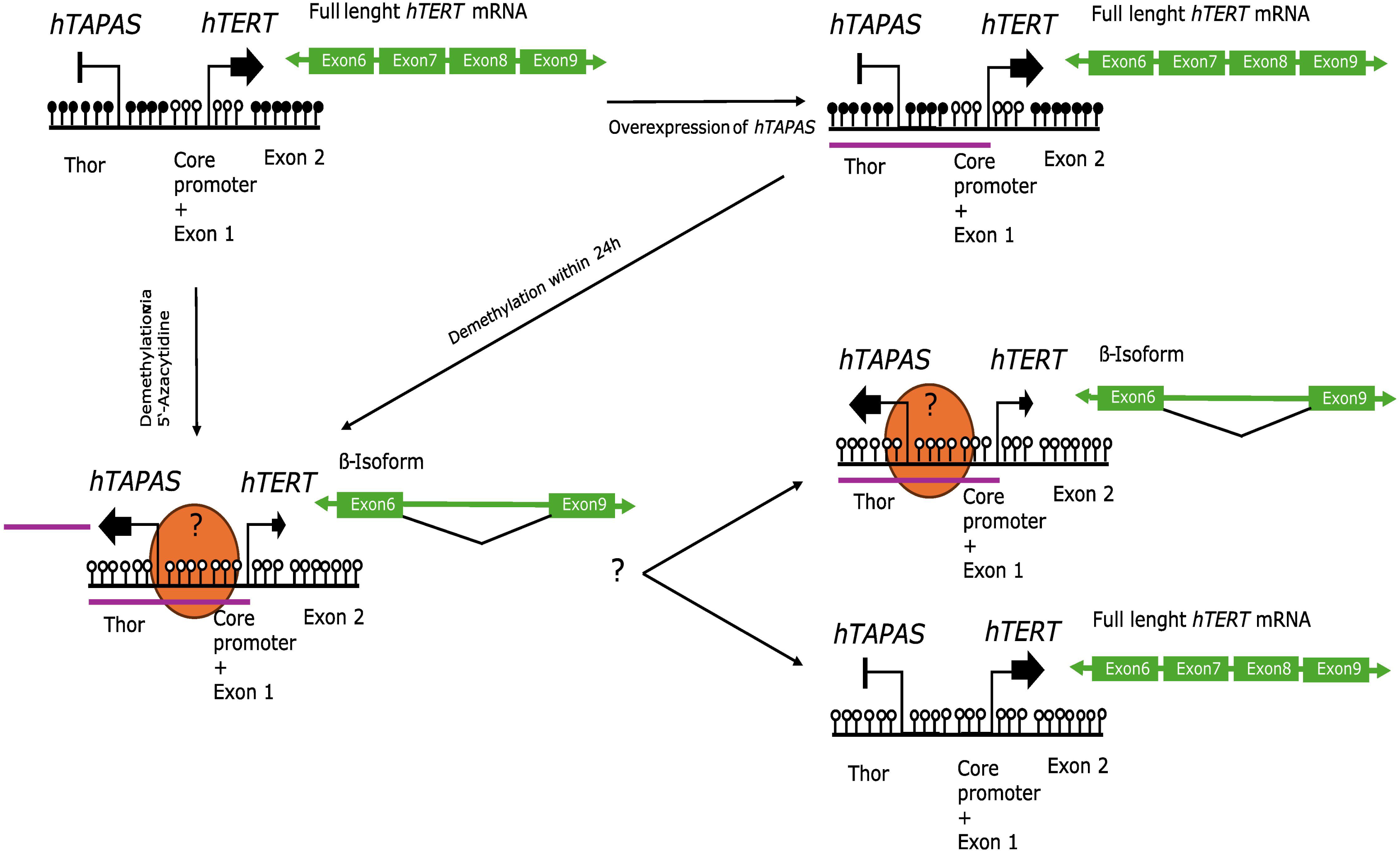
*hTAPAS* antisense transcription establishes a permissive epigenetic and splicing landscape at the *hTERT* locus. Antisense expression of *hTAPAS* maintains an unmethylated 5′ regulatory domain, including THOR and intron 2, promoting local chromatin accessibility and coordinating *hTERT* splice-site selection. Loss of *hTAPAS*, often driven by hypermethylation of its promoter in early tumorigenesis, shifts the locus toward a fully methylated state that favors production of full-length, catalytically active *hTERT* isoforms. DNA methylation and *hTAPAS* jointly influence the balance of full-length versus inactive β⁻/α⁻ *hTERT* transcripts, as illustrated by recapitulation of these effects through either *hTAPAS* overexpression or 5′-Azacytidine–mediated demethylation. Recovery of methylation and splicing patterns following withdrawal highlights the locus’s epigenetic memory. These findings indicate that antisense transcription and DNA methylation cooperate to regulate RNA polymerase II elongation and co-transcriptional splice-site recognition, providing a mechanistic link between epigenetic state and functional telomerase output with implications for telomerase reactivation in cancer.

## Material and Methods

### Cell lines and culture conditions

Human embryonic kidney 293T (HEK293T; ATCC® CRL-3216™), human embryonic lung fibroblast VA13 (WI-38 VA13 subline 2RA; ATCC® CCL-75.1™), human foreskin fibroblast HFF-1 (ATCC® SCRC-1041™), human lung fibroblast IMR-90 (ATCC® CCL-186™), and human foreskin fibroblast BJ (ATCC® CRL-2522™) cells were obtained from the American Type Culture Collection (ATCC, Manassas, VA, USA). Human B-cell precursor leukemia NALM-6 cells (DSMZ ACC 128) were obtained from the German Collection of Microorganisms and Cell Cultures (DSMZ, Braunschweig, Germany). Human induced pluripotent stem cells (iPSCs; Coriell GM26105, derived from participant huAA53E0 of the Personal Genome Project) were obtained from the Coriell Institute for Medical Research (Camden, NJ, USA).

#### HEK293T

HEK293T were maintained in Dulbecco’s Modified Eagle Medium (DMEM, high glucose; Gibco Thermo Fisher Scientific, Cat. No. 21969035) supplemented with 10% (v/v) heat-inactivated fetal bovine serum (FBS; PanBiotech, Cat. P30-3033) and 1% (v/v) penicillin–streptomycin (P/S; Gibco, Cat. No. 11548876). Cells were cultured at 37 °C in a humidified atmosphere containing 5% CO₂ and passaged at 70–80% confluence using 0.05% trypsin–EDTA (Gibco, Cat. No. 11528856). Reseeding was performed at a density of 2–4 × 10⁴ cells/cm².

#### VA13

VA13 were maintained in Eagle’s Minimum Essential Medium (EMEM, Gibco Thermo Fisher Scientific, Cat. No. 11440335) supplemented with 10% (v/v) heat-inactivated fetal bovine serum (FBS; PanBiotech, Cat. P30-3033) and 1% (v/v) penicillin–streptomycin (P/S; Gibco, Cat. No. 11548876). Cells were cultured at 37 °C in a humidified atmosphere containing 5% CO₂ and passaged at 70–80% confluence using 0.05% trypsin–EDTA (Gibco, Cat. No. 11528856). Reseeding was performed at a density of 2–4 × 10⁴ cells/cm².

#### HFF-1

HFF-1 cells were cultured in DMEM (high glucose) supplemented with 15% FBS and 1% P/S under standard culture conditions (37 °C, 5% CO₂). Cells were passaged with 0.05% trypsin–EDTA at 80–90% confluence and used for experiments before passage 15 to avoid replicative senescence.

#### IMR-90

IMR-90 cells were maintained in Eagle’s Minimum Essential Medium (EMEM; ATCC-formulated, Cat. No. 30-2003) supplemented with 10% FBS and 1% P/S. Cells were cultured at 37 °C with 5% CO₂ and split using 0.05% trypsin–EDTA when they reached 80–90% confluence. Experiments were performed with cells below passage 20 to minimize replicative senescence.

#### BJ

BJ fibroblasts were cultured in EMEM (ATCC-formulated, Cat. No. 30-2003) supplemented with 10% FBS and 1% P/S. Cells were maintained at 37 °C with 5% CO₂ and passaged using 0.05% trypsin–EDTA at 70–80% confluence. Cells were used for experiments before passage 15.

#### NALM-6

NALM-6 cells were maintained in RPMI 1640 medium (Gibco, Cat. No. 21875158) supplemented with 10% FBS and 1% P/S. Cultures were kept at densities between 2 × 10⁵ and 1 × 10⁶ cells/mL and split every 2–3 days by dilution with prewarmed medium.

Induced pluripotent stem cells (iPSCs)

Human iPSCs (HG002; Coriell GM26105) were maintained under feeder-free conditions in mTeSR™1 medium (STEMCELL Technologies, Cat. No. 85850) on six-well plates coated with hESC-qualified Matrigel® (Corning, Cat. No. 354277). Cultures were fed daily and maintained at 37 °C in a humidified atmosphere with 5% CO₂. Cells were passaged every 5–7 days using ReLeSR™ (STEMCELL Technologies, Cat. No. 05872) according to the manufacturer’s instructions. Only colonies with compact morphology and minimal spontaneous differentiation were selected for experiments.

### Cell Counting

Cells were counted using a Cellometer X1 automated cell counter (Nexcelom Bioscience LLC, Lawrence, Massachusetts, USA). For viability analysis, suspensions were mixed 1:1 with 0.4% trypan blue (Thermo Fisher Scientific, Cat. No 11538886) and 20 µL was loaded into disposable SD100 Cellometer counting chambers (CENIBRA, Belo Oriente, Minas Gerais, Brazil, Cat. No. CETHT4SD100014). Counts were performed using the manufacturer’s software with standard analysis settings, and results were reported as total cell concentration (cells/mL), viability (%), and live/dead counts. Each sample was analyzed in duplicate chambers, and values are presented as mean ± SD of at least three biological replicates.

### Transient Transfection

HEK293T and VA13 cells were seeded in 6-well plates at a density of 2 × 10⁵ cells/well 24 h prior to transfection to achieve 70–80% confluence at the time of transfection. Transfections were performed using Lipofectamine™ 3000 (Thermo Fisher Scientific, Cat. No. L3000-01) according to the manufacturer’s instructions. For each well, 2 µg plasmid DNA (pPlatTET-gRNA2 Addgene #82559, pcDNA3.1(+) eGFP Addgene #129020 or pCDNA-3xHA-*hTERT* Addgene #51637) was diluted in 125 µL Opti-MEM™ I Reduced Serum Medium (Gibco, Cat. No. 15392402) with 5 µL P3000™ reagent. Separately, 7.5 µL Lipofectamine™ 3000 was diluted in 125 µL Opti-MEM™. The diluted DNA and Lipofectamine™ 3000 were combined, incubated for 10 min at room temperature, and added dropwise to the cells. After 6 h, the medium was replaced with fresh complete growth medium. Cells were harvested for analysis 24–144 h post-transfection. pcDNA3.1(+) eGFP was a gift from Jeremy Wilusz (Addgene plasmid # 129020 ; http://n2t.net/addgene:129020 ; RRID:Addgene_129020)[33].

### Generation of *hTAPAS* and *hSAPAT* expression constructs

Cloning of *hTAPAS* and *hSAPAT* was carried out using PCR primers (Integrated DNA Technologies, IDT) containing AscI (NEB, Cat. No. R0558S) and AgeI (NEB, Cat. No. R3552S) restriction sites. The expression vector pCDNA-3xHA-*hTERT* (Addgene plasmid #51637; RRID:Addgene_51637), a gift from Steven Artandi[34], was used as the backbone. One microgram of the plasmid was digested with AscI and AgeI at 37 °C for 1 h to excise the *hTERT* coding region and generate compatible ends for insertion of the lncRNA or antisense transcript. The digested PCR products and linearized plasmids were ligated overnight at 14 °C using T4 DNA ligase (NEB). Ligation products were transformed into One Shot™ TOP10 Chemically Competent E. coli (Thermo Fisher Scientific, Cat. No. 10368022), and transformants were selected on kanamycin-containing agar. Positive colonies were screened by colony PCR and verified by Sanger sequencing using the M13 Forward primer. Plasmids containing verified inserts were purified using the QIAprep Spin Miniprep Kit (Qiagen, Cat. No. 27106).

### Cell Sorting

Cell sorting was performed on the Invitrogen Bigfoot Spectral Cell Sorter (Thermo Fisher Scientific) or the BD FACSAria III SORP (BD Biosciences) using a 100 µm nozzle and 20 psi setup and Purity Sort Precision Mode. HEK293T and VA13 cells were prepared in sterile phosphate-buffered saline (PBS) supplemented with 2% fetal bovine serum (FBS) and filtered through 40 µm nylon mesh to remove aggregates prior to sorting. Cell suspensions were kept on ice until acquisition. Cells were gated based on FSC-A vs SSC-A properties. Doublets were excluded using SSC-A vs. SSC-H. DAPI was included for live/dead discrimination (0.5 µg/ml final concentration). The GFP-positive population was defined using an untransfected cell line for the GFP-cutoff. Sorted populations were collected into 15ml Falcon tubes containing culture medium supplemented with 10% FBS

### 5’-Azacytidine Treatment

HEK293T and VA13 cells were seeded at 60–70% confluence and treated with 1 µM 5′-Azacytidine (Sigma-Aldrich, St. Louis, MO, USA, Cat. No. A3656-5MG) freshly prepared in sterile DMSO. The treatment was carried out in complete growth medium for 120 h, with daily replacement of medium containing freshly diluted drug. Control cells received an equivalent volume of DMSO.

### High–Molecular-Weight DNA Isolation

Genomic DNA was isolated using the Blood & Cell Culture DNA Mini Kit (25) (Qiagen, Hilden, Germany; Cat. No. 13343) with modifications to preserve DNA integrity. All pipetting of cell lysates and DNA solutions was performed using wide-bore, low-retention tips (20 µL/gauge) to minimize mechanical shearing. Lysis was performed by gentle inversion mixing, avoiding vortexing. DNA precipitation and washing followed the manufacturer’s instructions. DNA was eluted in TE buffer prewarmed to 50 °C.

### RNA Preparation, cDNA Synthesis, and Real-Time PCR

Total RNA was isolated from cultured cells using the RNeasy Mini Kit (Qiagen; Cat. No. 74104) according to the manufacturer’s instructions. Briefly, cells were lysed directly in RLT buffer containing β-mercaptoethanol, and lysates were homogenized using QIAshredder spin columns to reduce viscosity. RNA was bound to the silica membrane, washed, and eluted in RNase-free water. RNA concentration and purity were assessed using a NanoDrop™ spectrophotometer (Thermo Fisher), and integrity was confirmed by agarose gel electrophoresis.

For complementary DNA (cDNA) synthesis, 1 µg of total RNA was reverse-transcribed using the SuperScript™ VILO cDNA Synthesis Kit (Thermo Fisher Scientific; Cat. No. 11754250) following the manufacturer’s protocol. Briefly, RNA was mixed with the VILO Reaction Mix and VILO Enzyme Mix, incubated at 25 °C for 10 min, followed by 42 °C for 60 min, and then inactivated at 85 °C for 5 min. The resulting cDNA was stored at −20 °C.

Real-time PCR was performed using PerfeCTa® SYBR® Green FastMix™ with Low-ROX reference dye (VWR, Radnor, PA, USA; Cat. No. 733-1390) on an Applied Biosystems QuantStudio 5 Real-Time PCR System (Thermo Fisher Scientific) using MicroAmp™ Optical 384-Well Reaction Plates (Fisher Scientific; Cat. No. 10005724). Reactions were set up in a total volume of 10 µL, containing 1× SYBR Green FastMix, 0.3 µM of each primer, and 2 µL of cDNA template. Plates were sealed and centrifuged briefly to remove bubbles. qPCR cycling conditions were: initial denaturation at 95 °C for 3 min, followed by 40 cycles of 95 °C for 10 s and 60 °C for 30 s. Melt curve analysis was included to verify amplification specificity. Relative *hTERT* and *hTAPAS* expression was calculated using the ΔΔCt method with *GAPDH* as housekeeping gene reference. Primer sequences are given in supplementary table 1.

### Isoform detection PCR

Detection of *hTERT* splice isoforms encompassing exons 5–9 was performed using a nested PCR approach, whereas isoforms spanning exons 2–3 were analyzed by conventional PCR. All reactions were carried out using NEB Taq DNA Polymerase (New England Biolabs; Cat. No. M0273X) in accordance with the manufacturer’s instructions. The first round of nested PCR comprised 30 cycles of denaturation at 95 °C for 30 s, annealing at 54 °C for 40 s, and extension at 72 °C for 25 s. The second (nested) amplification was conducted for 40 cycles under identical denaturation and extension conditions, with an increased annealing temperature of 59 °C for 40 s. Conventional PCR for exons 2–3 was performed for 30 cycles with denaturation at 95 °C for 30 s, annealing at 60 °C for 40 s, and extension at 72 °C for 25 s. Amplification products were resolved by agarose gel electrophoresis to verify the expected amplicon sizes. Primer sequences are provided in supplementary Table 2.

### CRISPR-Cas9 Enrichment and Oxford Nanopore Sequencing

CRISPR-Cas9 targeted enrichment was performed as described in [35]. Cas9 ribonucleoprotein (RNP) complexes were prepared by annealing pooled crRNA probes with Alt-R® CRISPR-Cas9 tracrRNA, 5 nmol (IDT, Coralville, IA, USA; Cat. No 1072532) in Duplex Buffer (10 µl total), followed by assembly with Alt-R® S.p. HiFi Cas9 Nuclease V3, 100 µg (IDT; Cat. No 1081060) and rCutSmart Buffer. Genomic DNA was dephosphorylated using Quick CIP (NEB; Cat. No M0525S) at 37°C for 15 min, then cleaved and dA-tailed in the presence of Cas9 RNPs, NEB Taq DNA Polymerase and dATP. Libraries were barcoded using ONT Native Barcodes (Oxford Nanopore Technologies, Oxford, UK; Cat. No SQK-NBD114.24) and ligated with NEB Next Quick Ligation reagents (NEB; Cat. No M2200S). AMPure XP beads (Beckman Coulter, Brea, CA, USA; Cat. No A63881) were used for post-ligation clean-up, and DNA was eluted in nuclease-free water at 37°C, followed by overnight incubation at 4°C. Flow cells were primed with Flow Cell Flush (FCF) and Flow Cell Tether (FCT) containing BSA, and libraries were loaded via the SpotON port for sequencing on MinION devices using MinKNOW software. crRNA Probe sequences are given in supplementary table 3+4. Raw Oxford Nanopore POD5 files were basecalled using Dorado (v0.8.1), using the super high-accuracy “sup” model with modified base detection for 5mC and 5hmC. Although a newer version of Dorado is available, v0.8.1 was used to ensure consistency across all processed datasets. Basecalling was performed on a CUDA-enabled GPU (cuda:0) with recursive processing of input files, generating BAM outputs aligned to the T2T-CHM13v2.0 human reference genome.

Reads were demultiplexed using Dorado with EXP-NBD114 and EXP-NBD104 kit with BAM sorting enabled. Demultiplexed BAM files were subsequently aligned to the T2T-CHM13v2.0 reference using the Dorado aligner. Aligned reads were sorted and indexed using Samtools (v1.20). DNA methylation profiles in CpG context were extracted using Modkit pileup with reference-guided processing. Final methylation outputs were generated in BED format and visualised using custom Python scripts. Coverage ranged from 4× to 300×, with an average depth of 32× across cell lines and enriched genomic loci (Supplementary Figures 2 and 3).

## Declarations

### Ethics approval and consent to participate

Not applicable.

### Consent for publication

Not applicable.

### Availability of data and materials

All data generated or analysed during this study are included in this published article [and its supplementary information files].

### Competing interests

The authors declare that they have no competing interests.

### Funding

Not applicable.

### Authors’ contributions

L.E. conceived the study and carried out cell culture, nanopore sequencing, qRT-PCR, and splicing experiments. He was also responsible for data curation, visualization, and drafting as well as revising the manuscript. V.S.J.P. conducted the bioinformatic analyses and contributed to data visualization. A.G.L. and J.A.D. performed cell culture and splicing experiments. C.K. developed the methodology for nanopore multiplexing.

## Acknowledgements

The authors would like to express their sincere gratitude to Prof. Dr. Peter Baumann for his generous support in providing the funding and research infrastructure that made this project possible. They further thank the IMB Flow Cytometry Core Facility for their expert assistance. Cell sorting was performed using the BD FACS Aria™ III SORP (DFG Project No. 210144599) and the Invitrogen Bigfoot Spectral Cell Sorter (DFG Project No. 511658729). The authors are grateful to Prof. Dr. Simeon Santourlidis for revising the paper.

## Notes

### Competing Interest Statement

The authors have declared no competing interest.

