## supplementary figure for "Epigenetic–splicing regulation of *hTERT* mediated by *hTAPAS*"

### Supplementary figure 1: Sanger seq result for PCR product of hTERT ex2-3 and 5-9.

```
Ex_2_3      ------------------------------------------------------------
Seq13       CCCGACCTACTCTATAGGGCGATTGGGCCCTCTAGATGCATGCTCGAGCGGCCGCCAGTG

Ex_2_3      --------------------------GAGGAGGAGGACACAGACCCCCGTCGCCTGGTGC
Seq13       TGATGGATATCTGCAGAATTCGCCCTTAGGAGGAGGACACAGACCCCCGTCGCCTGGTGC
                                       *********************************

Ex_2_3      AGCTGCTCCGCCAGCACAGCAGCCCCTGGCAGGTGTACGGCTTCGTGCGGGCCTGCCTGC
Seq13       AGCTGCTCCGCCAGCACAGCAGCCCCTGGCAGGTGTACGGCTTCGTGCGGGCCTGCCTGC
            ************************************************************

Ex_2_3      GCCGGCTGGTGCCCCCAGGCCTCTGGGGCTCCAGGCACAACGAACGCCGCTTCCTCAGGA
Seq13       GCCGGCTGGTGCCCCCAGGCCTCTGGGGCTCCAGGCACAACGAACGCCGCTTCCTCAGGA
            ************************************************************

Ex_2_3      ACACCAAGAAGTTCATCTCCCTGGGGAAGCATGCCAAGCTCTCGCTGCAGGAGCTGACGT
Seq13       ACACCAAGAAGTTCATCTCCCTGGGGAAGCATGCCAAGCTCTCGCTGCAGGAGCTGACGT
            ************************************************************

Ex_2_3      GGAAGATGAGCGTGCGGGACTGCGCTTGGCTGCGCAGGAGCCCAGGGGTTGGCTGTGTTC
Seq13       GGAAGATGAGCGTGCGGGACTGCGCTTGGCTGCGCAGGAGCCCAGGGGTTGGCTGTGTTC
            ************************************************************

Ex_2_3      CGGCCGCAGAGCACCGTCTGCGTGAGGAGATCCTGGCCAAGTTCCTGCACTGGCTGATGA
Seq13       CGGCCGCAGAGCACCGTCTGCGTGAGGAGATCCTGGCCAAGTTCCTGCACTGGCTGATGA
            ************************************************************

Ex_2_3      GTGTGTACGTCGTCGAGCTGCTCAGGTCTTTCTTTTATGTCACGGAG
Seq13       GTGTGTACGTCGTCGAGCTGCTCAGGTCTTTCTTTTATGTCACGGAG
            ***********************************************
```

```
ß-deletion     GGACAGGCTCACGGAGGTCATCGCCAGCATCATCAAACCCCAGAACACGTACTGCGTGCG
ß-deletion     GGACAGGCTCACGGAGGTCATCGCCAGCATCATCAAACCCCAGAACACGTACTGCGTGCG
Full length    GGACAGGCTCACGGAGGTCATCGCCAGCATCATCAAACCCCAGAACACGTACTGCGTGCG
               ************************************************************

ß-deletion     TCGGTATGCCGTGGTCCAGAAGGCCGCCCATGGGCACGTCCGCAAGGCCTTCAAGAGCCA
ß-deletion     TCGGTATGCCGTGGTCCAGAAGGCCGCCCATGGGCACGTCCGCAAGGCCTTCAAGAGCCA
Full length    TCGGTATGCCGTGGTCCAGAAGGCCGCCCATGGGCACGTCCGCAAGGCCTTCAAGAGCCA
               ************************************************************

ß-deletion     C-----------------------------------------------------------
ß-deletion     C-----------------------------------------------------------
Full length    CGTCTCTACCTTGACAGACCTCCAGCCGTACATGCGACAGTTCGTGGCTCACCTGCAGGA
               *

ß-deletion     ------------------------------------------------------------
ß-deletion     ------------------------------------------------------------
Full length    GACCAGCCCGCTGAGGGATGCCGTCGTCATCGAGCAGAGCTCCTCCCTGAATGAGGCCAG

ß-deletion     ------------------------------------------------------------
ß-deletion     ------------------------------------------------------------
Full length    CAGTGGCCTCTTCGACGTCTTCCTACGCTTCATGTGCCACCACGCCGTGCGCATCAGGGG

ß-deletion     ---GTCCTACGTCCAGTGCCAGGGGATCCCGCAGGGCTCCATCCTCTCCACGCTGCTCTG
ß-deletion     ---GTCCTACGTCCAGTGCCAGGGGATCCCGCAGGGCTCCATCCTCTCCACGCTGCTCTG
Full length    CAAGTCCTACGTCCAGTGCCAGGGGATCCCGCAGGGCTCCATCCTCTCCACGCTGCTCTG
                  *********************************************************
```

Supplementary Figure 2: Intron 6-8 methylation

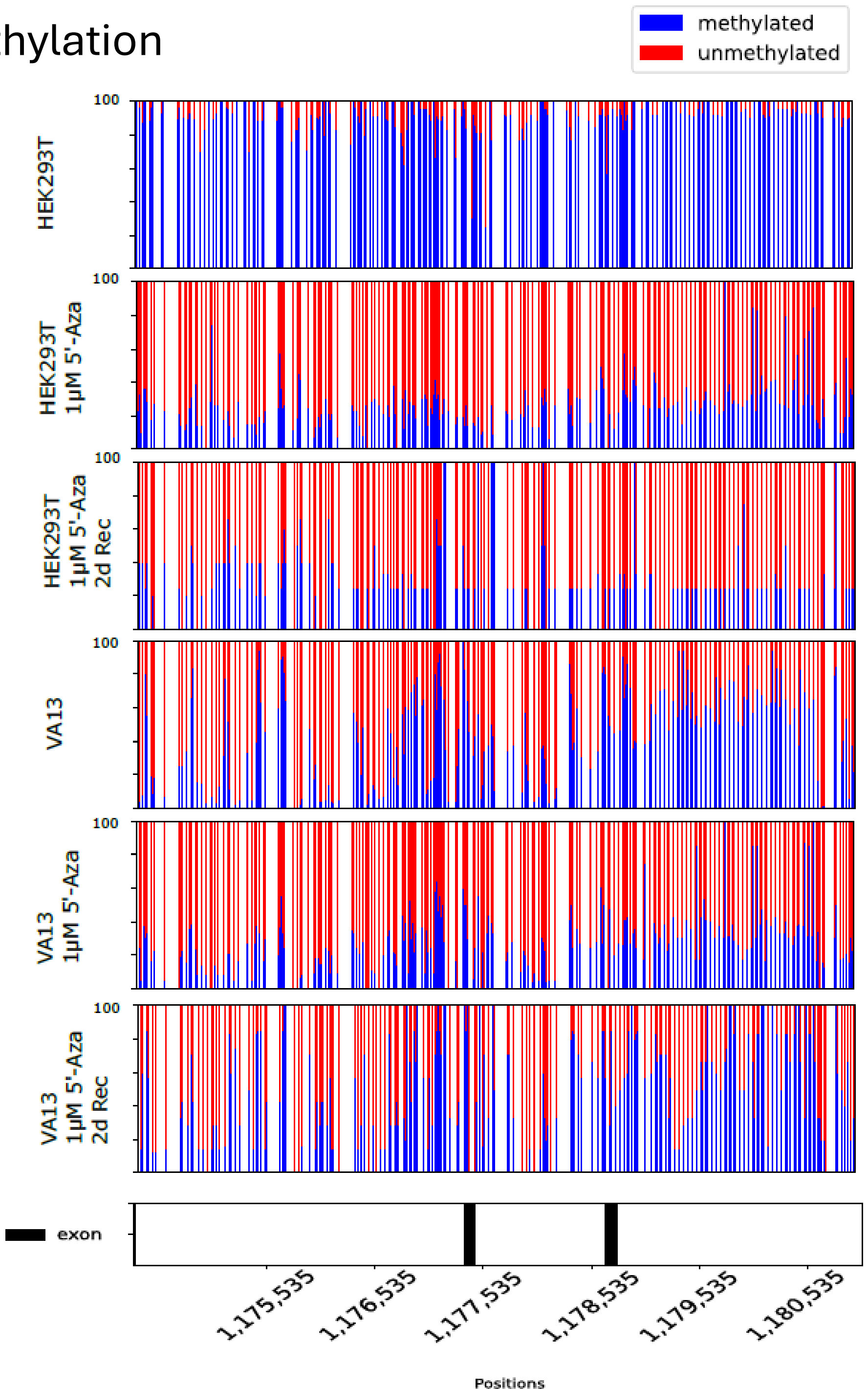

### Supplementary figure 2: Intron 6-8 methylation and coverage plots for HEK293T and VA13 cells after 5’-Azacytidine treatment

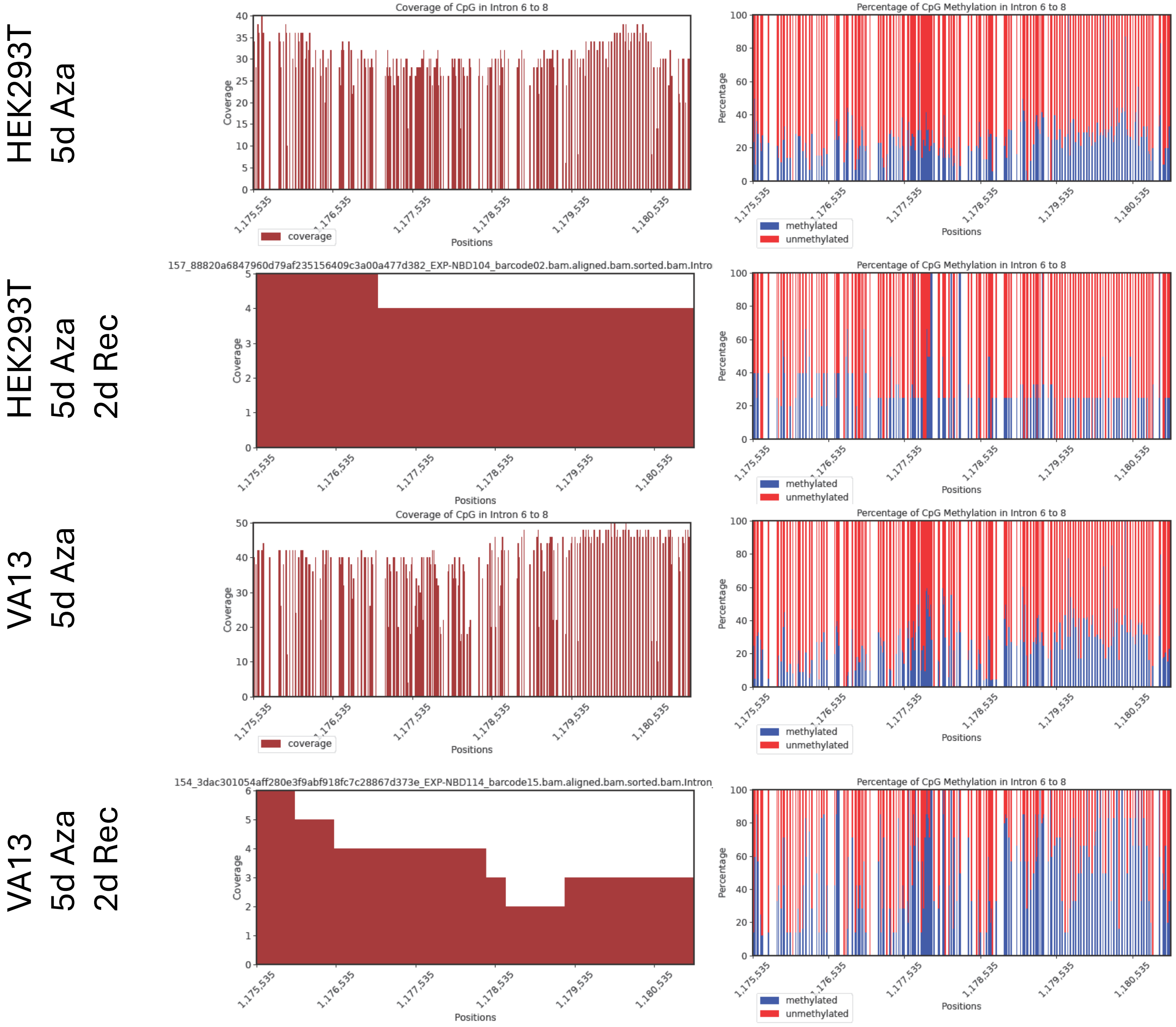

### Supplementary figure 3: Coverage Plots.

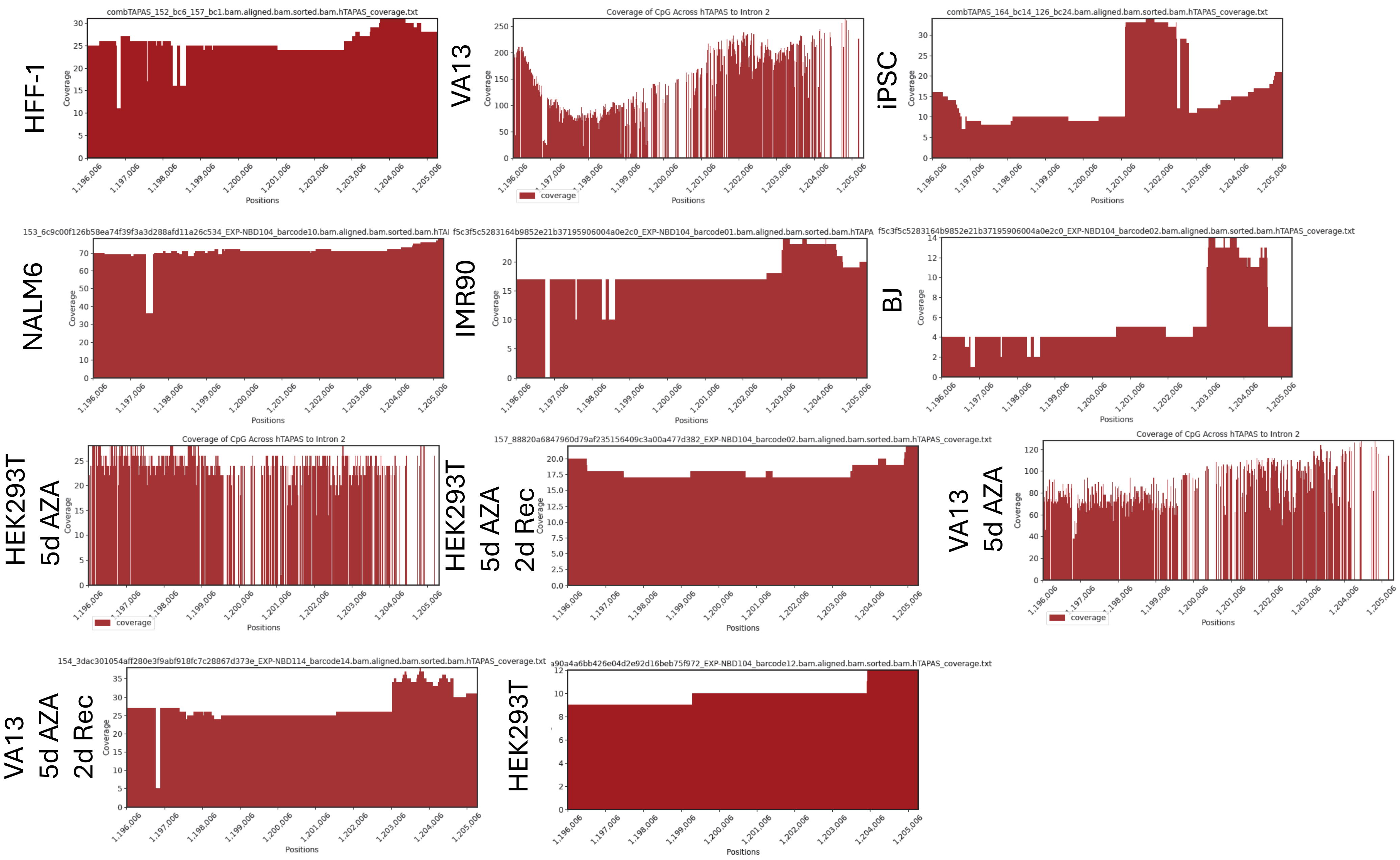

### Supplementary figure 3: Coverage Plots.

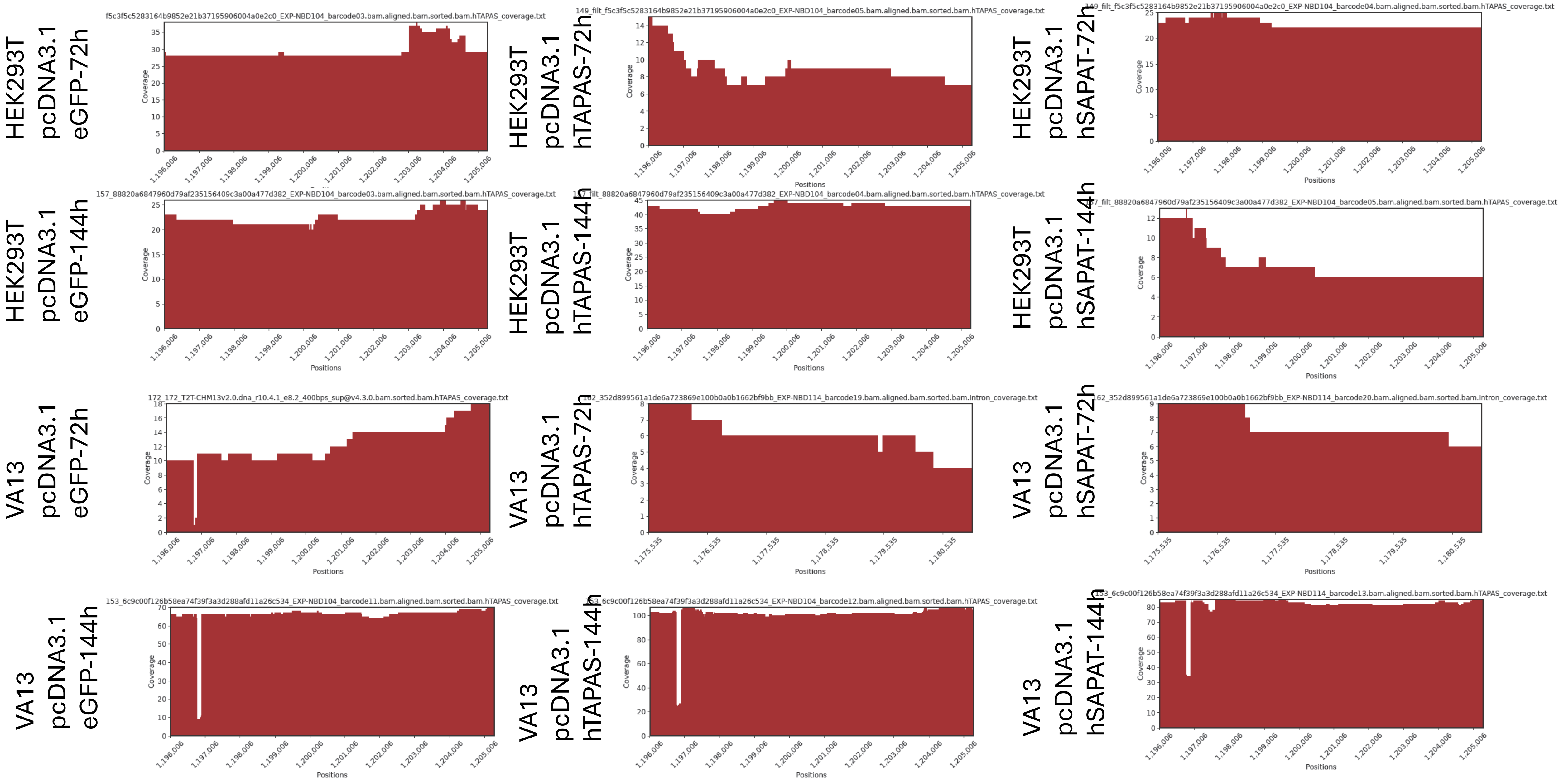

Supplementary figure 3: Coverage Plots intron 6-8.

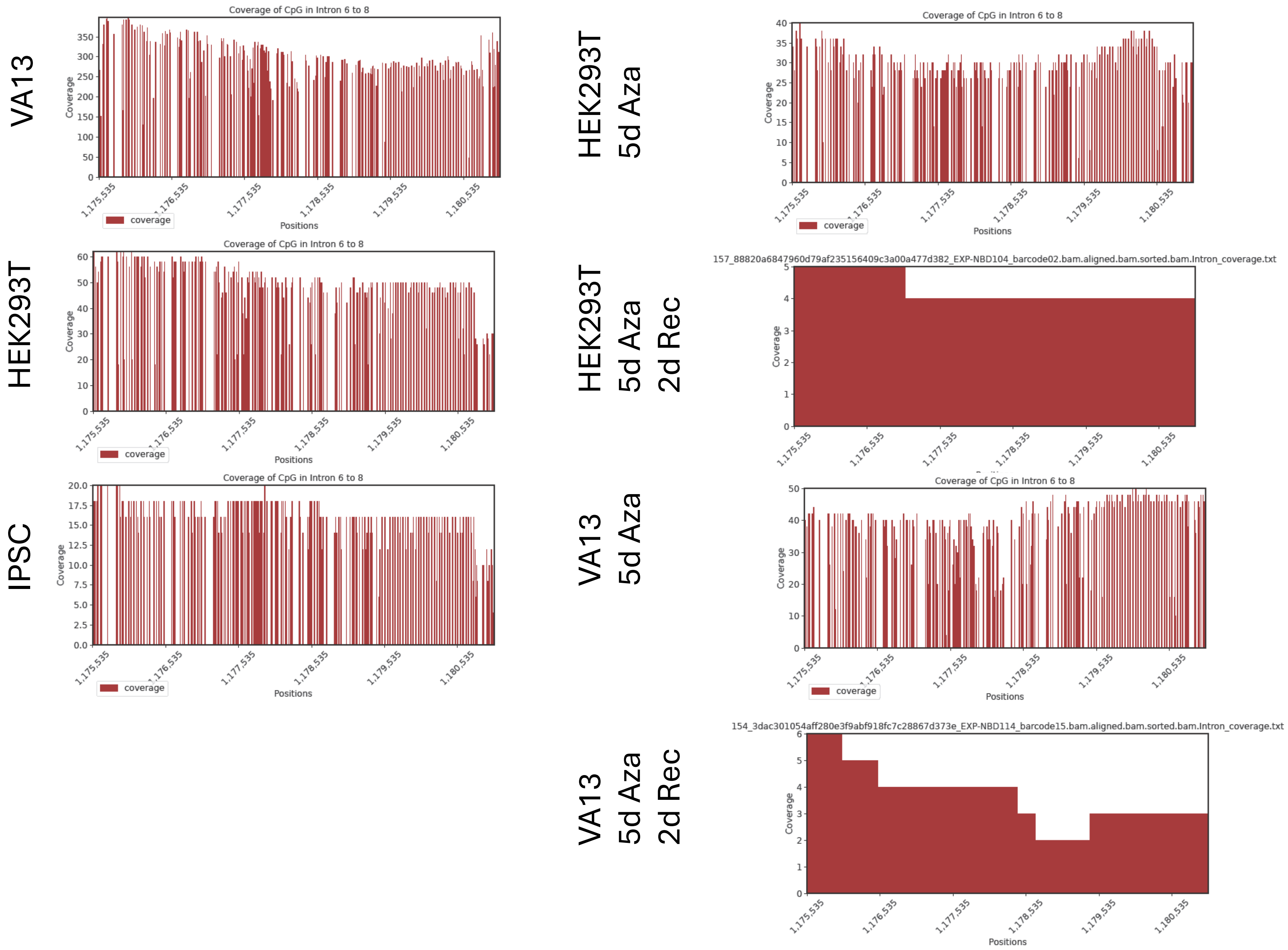

#### Supplementary figure 4: FACS sorted GFP positive VA13 and HEK293T cells

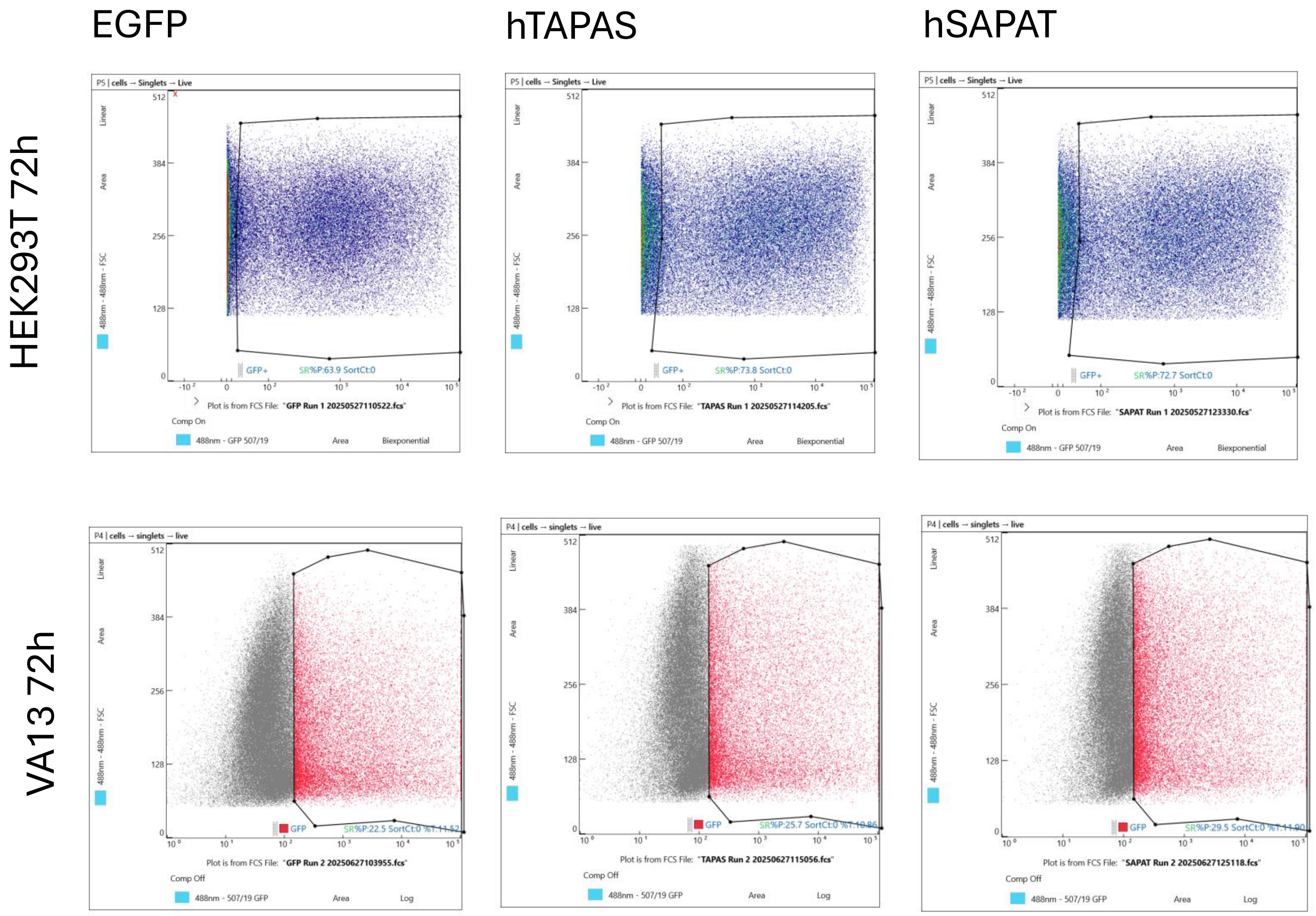

Supplementary figure 5: Fluorescent microscope images of GFP positive VA13 and HEK293T cells

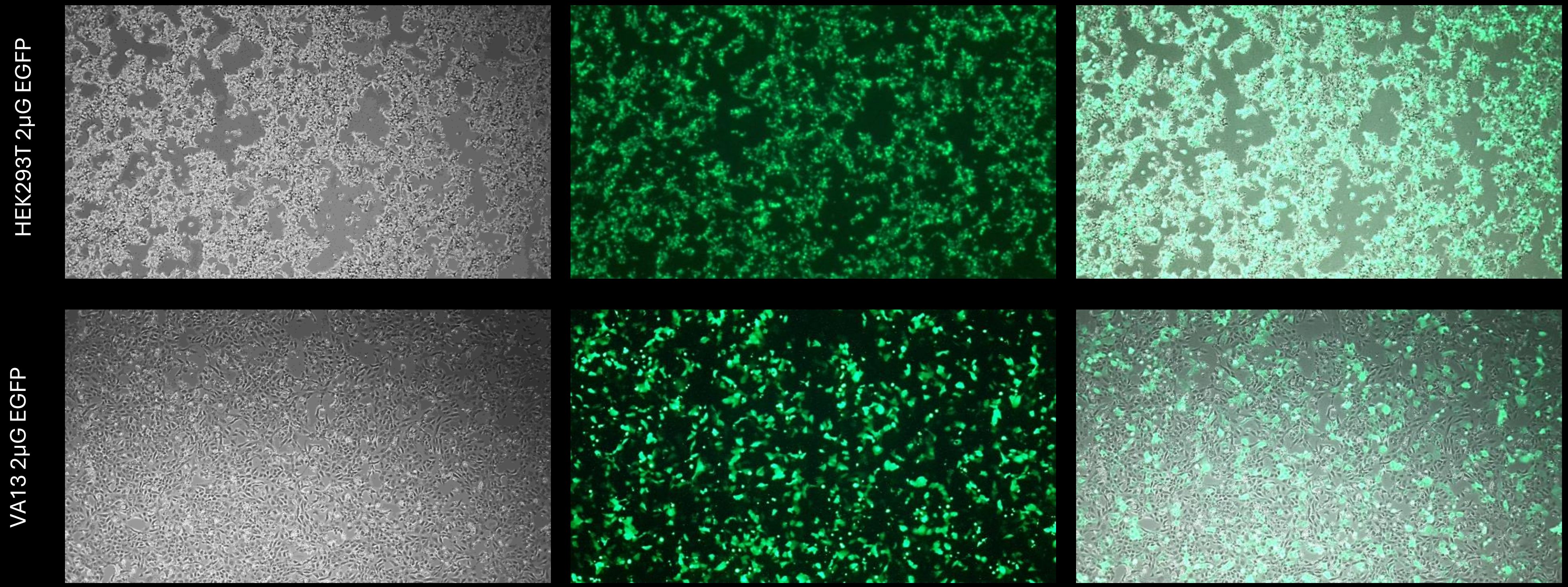

### Supplementary figure 6: qRT-PCR results for the expression levels of the lncRNA *hTAPAS*

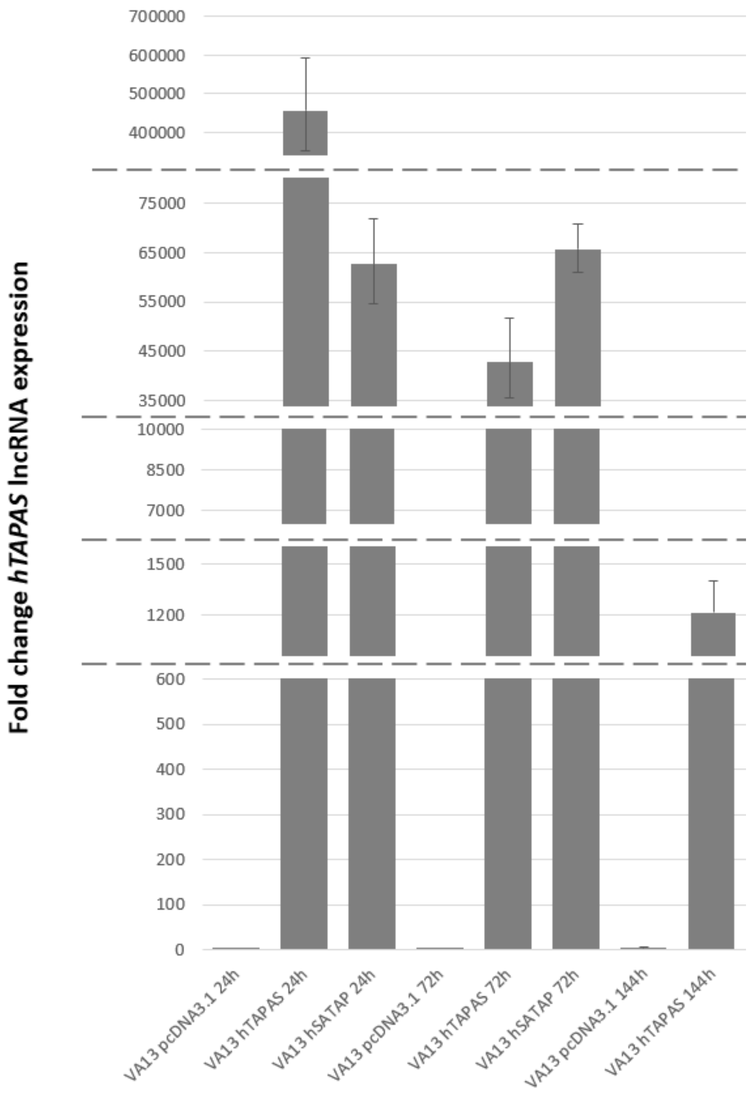


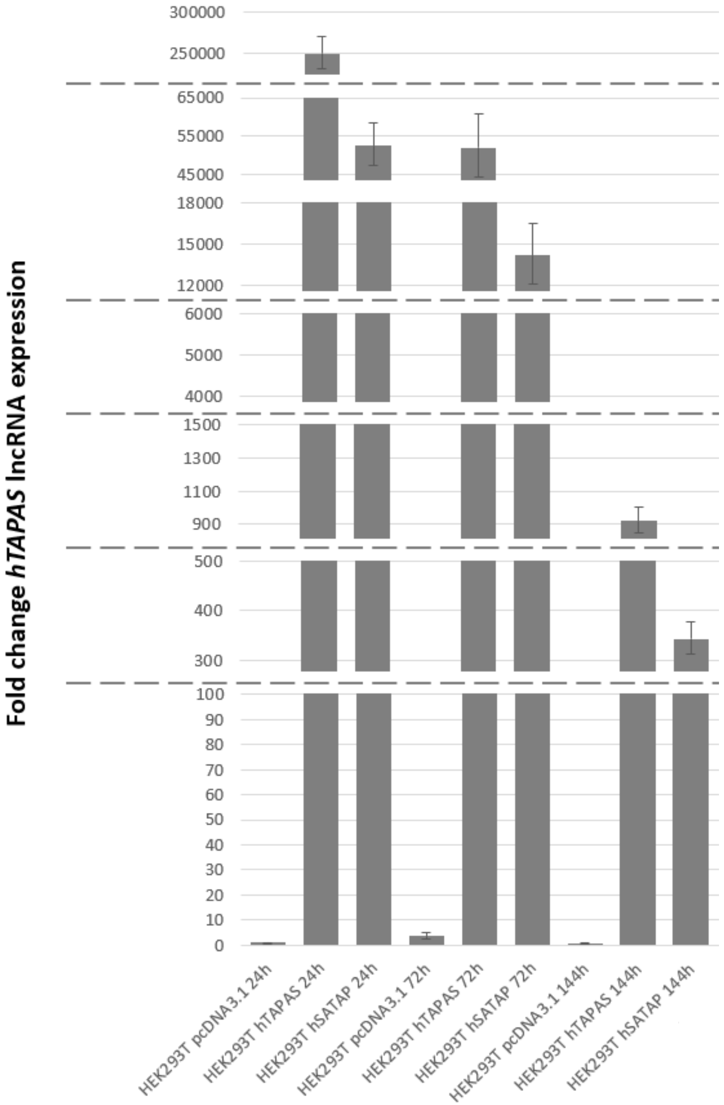

### Supplementary figure 7: Bisulfite sequencing for hTERT -482 to -696.

120h HEK293T

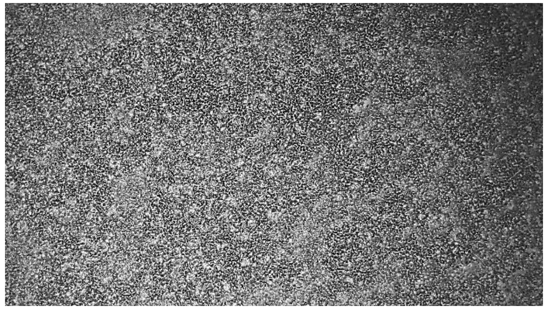

| CpG position | 25 | 27 | 53 | 63 | 65 | 71 | 87 | 93 | 99 | 109 | 111 | 117 | 119 | 125 | 139 |
|---|---|---|---|---|---|---|---|---|---|---|---|---|---|---|---|
| Me-CpG | 10/10 100.0% | 9/10 90.0% | 10/10 100.0% | 10/10 100.0% | 9/10 90.0% | 9/10 90.0% | 10/10 100.0% | 10/10 100.0% | 10/10 100.0% | 10/10 100.0% | 10/10 100.0% | 10/10 100.0% | 10/10 100.0% | 10/10 100.0% | 9/10 90.0% |
| **CpG position** | **143** | **150** | **159** | **166** | **169** | **174** | **176** | **180** | **206** | **219** | **231** | **234** | **238** | | **Total** |
| Me-CpG | 10/10 100.0% | 10/10 100.0% | 10/10 100.0% | 9/10 90.0% | 10/10 100.0% | 10/10 100.0% | 10/10 100.0% | 10/10 100.0% | 10/10 100.0% | 10/10 100.0% | 10/10 100.0% | 10/10 100.0% | 10/10 100.0% | | 275/280 98.2% |

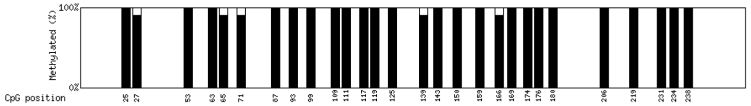


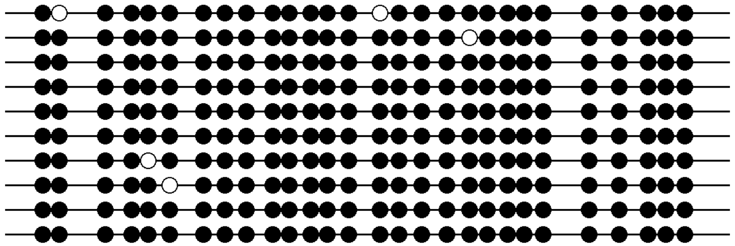

120h HEK293T treated with 1μM Aza

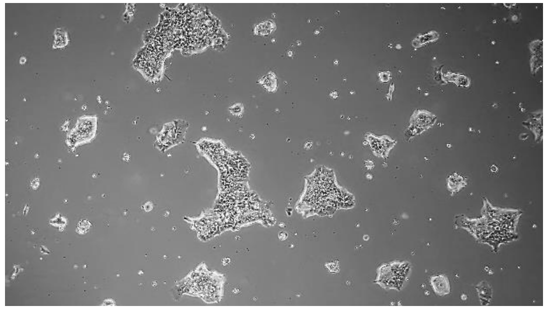

| CpG position | 18 | 20 | 46 | 56 | 58 | 64 | 80 | 86 | 92 | 102 | 104 | 110 | 112 | 118 | 132 |
|---|---|---|---|---|---|---|---|---|---|---|---|---|---|---|---|
| Me-CpG | 5/10 50.0% | 7/10 70.0% | 4/10 40.0% | 4/10 40.0% | 3/10 30.0% | 4/10 40.0% | 5/10 50.0% | 5/10 50.0% | 6/10 60.0% | 8/10 80.0% | 6/10 60.0% | 7/10 70.0% | 7/10 70.0% | 8/10 80.0% | 7/10 70.0% |
| **CpG position** | **136** | **143** | **152** | **159** | **162** | **167** | **169** | **173** | **199** | **212** | **224** | **227** | **231** | | **Total** |
| Me-CpG | 8/10 80.0% | 7/10 70.0% | 9/10 90.0% | 8/10 80.0% | 9/10 90.0% | 10/10 100.0% | 9/10 90.0% | 9/10 90.0% | 9/10 90.0% | 9/10 90.0% | 9/10 90.0% | 10/10 100.0% | 10/10 100.0% | | 202/280 72.1% |

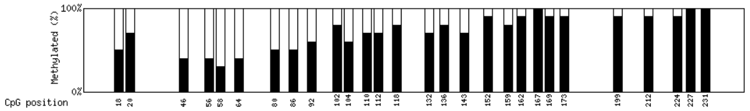


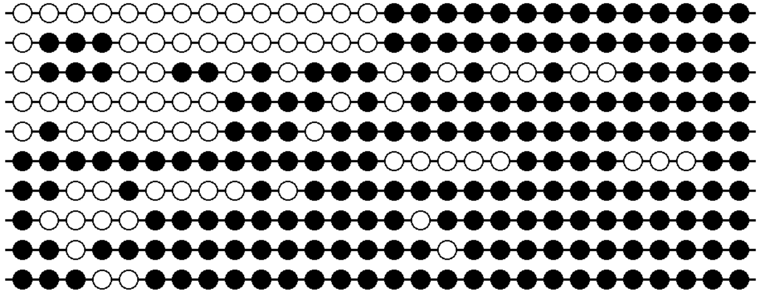
