## supplementary tables for "Epigenetic–splicing regulation of *hTERT* mediated by *hTAPAS*"

Table 1: Real Time PCR Primers

| Primer | Sequence | TM |
|---|---|---|
| hTAPAS s | 5‘-tgtagctgaggtcggcaaac-3‘ | 60°C |
| hTAPAS as | 5‘-ggtgcgaggcctgttcaaat-3‘ | |
| hTERT s | 5‘-cggaagagtgtctggagcaa-3‘ | 60°C |
| hTERT as | 5‘-ggatgaagcggagtctgga-3‘ | |
| GAPDH s | 5‘-acatcgctcagacaccatg-3‘ | 60°C |
| GAPDH as | 5‘-tgtagttgaggtcaatgaaggg-3‘ | |

Table 2: Isoform detection PCR

| Primer | Sequence | TM |
|---|---|---|
| hTERT e3-13 s | 5‘-gctgctcaggtctttcttttat-3‘ | 54°C |
| hTERT e3-13 as | 5‘-ggaggatcttgtagatgttggt-3‘ | |
| hTERT e5-9 s | 5‘-gcctgagctgtactttgtcaa-3‘ | 59°C |
| hTERT e5-9 as | 5‘-cgcaaacagcttgttctccatgtc-3‘ | |
| hTERT e2-3 s | 5‘-gaggaggaggacacagacccc-3‘ | 63°C |
| hTERT e2-3 as | 5‘-ctccgtgacataaaagaaagacctg-3‘ | |

Table 3: crRNA probes to enrich for Chr. 5: 1,196,006–1,205,206

| Probe | Sequence |
|---|---|
| 1 | 5’- GTGACCACCTGTTATCCCAT-3‘ |
| 2 | 5‘-TCCATGAACTCCTTACCAGT-3‘ |
| 3 | 5’- GGACGTCAATCCATGTGAGG-3‘ |
| 4 | 5’- AGCGTTGCCACCCACCCAAG-3‘ |

Table 4: crRNA probes to enrich for Chr. 5: 1,174,035-1,180,535

| Probe | Sequence |
|---|---|
| 1 | 5’-gggtgtagaccccatgcaag-3’ |
| 2 | 5’-tgagttgaaccacattagg-3’ |
| 3 | 5’-aatatatcaacactgacgag-3’ |
| 4 | 5’-tttgatgcctcacaagctcg-3’ |
